# A “Human-in-the-Loop” Approach for Semi-automated Image Restoration in Electron Microscopy

**DOI:** 10.1101/644146

**Authors:** Joris Roels, Frank Vernaillen, Anna Kremer, Amanda Gonçalves, Jan Aelterman, Hiêp Q. Luong, Bart Goossens, Wilfried Philips, Saskia Lippens, Yvan Saeys

## Abstract

The recent advent of 3D in Electron Microscopy (EM) has allowed for detection of detailed sub-cellular nanometer resolution structures. While being a scientific breakthrough, this has also caused an explosion in dataset size, necessitating the development of automated workflows. Automated workflows typically benefit reproducibility and throughput compared to manual analysis. The risk of automation is that it ignores the expertise of the microscopy user that comes with manual analysis. To mitigate this risk, this paper presents a hybrid paradigm. We propose a ‘human-in-the-loop’ (HITL) approach that combines expert microscopy knowledge with the power of large-scale parallel computing to improve EM image quality through advanced image restoration algorithms. An interactive graphical user interface, publicly available as an ImageJ plugin, was developed to allow biologists to use our framework in an intuitive and user-friendly fashion. We show that this plugin improves visualization of EM ultrastructure and subsequent (semi-)automated segmentation and image analysis.

## Introduction

The field of three-dimensional electron microscopy (3D EM) covers several technologies that unveil a sample at nanometer (nm) resolution. Although 3D EM has been performed for several decades, the development of Serial Block Face (SBF) Scanning EM (SEM) has made 3D EM more easily available for large scale imaging of biological samples^1^. SBF-SEM repetitively acquires a 2D SEM image from the sample surface and sections the top of the sample using a diamond knife ultramicrotome^2,3^, which results in a 3D stack. Another block face imaging technique, where ‘slice and view’ is used, is Focused Ion Beam (FIB) SEM, where the block face is removed by FIB milling. While both SBF-SEM and FIB-SEM have the potential to generate images at (3 to 5) nm lateral resolution, the FIB milling is more precise than the mechanical SBF-SEM slicing, resulting in a maximal axial resolution of 5 nm and 20 nm, respectively^1, 4, 5^.

Over the past years, there have been staggering advances in the use of these techniques in life science research due to their throughput in collecting 3D EM images^6–10^. The strength of EM is not only its resolution, but also that all subcellular information is present in one image. Additionally, the fundamental advantage of 3D information at high resolution is that the researcher can generate a comprehensive view of a complete cell or tissue.

Although datasets are generated in a relatively short time, data processing is a major bottleneck in the workflow. Data interpretation is very challenging due to the density of information, which makes distinguishing individual features and their 3D rendering less than trivial. For that reason, segmentation — the process of indicating the boundaries of a biological object — is an essential step in data interpretation and analysis of 3D EM images, although it is largely responsible for the bottleneck in SEM data analysis. In cases where a targeted object has been specifically stained and the pixel intensity stands out from the rest of the image, basic techniques such as intensity thresholding can be applied^11^. However, thresholding fails in most cases, and segmentation is done manually^12^, which has several issues. Firstly, such a process is very labor-intensive, making it slow and costly. Secondly, reproducibility is an issue since humans tends to be subjective and biased, and different annotators are likely to produce different segmentations. Lastly, segmentation quality depends on high-quality data, prompting the introduction of quality control.

Automated analysis is key to solving the bottleneck in the entire 3D EM workflow. Current advances in large-scale computing and computer vision^13–15^ can significantly speed up the whole automated image analysis process and make it more cost efficient, potentially leading to challenging and successful research projects that were never possible before^16^. Furthermore, automated methods typically have a formal mathematical formulation and thus the potential to be less biased and more reproducible. Recent automated analysis frameworks are all focused on the segmentation process as such, by either facilitating the manual tracing^17, 18^ or providing algorithms that allow for interactive learning^19^. As data quality control is omitted, these methods often underperform when challenged with image artifacts such as blur and noise. In practice, this is fairly common in volume EM considering the physical and theoretical issues, including imperfect lenses, thermal heating, diffraction and electron counting errors^20^. Moreover, these artifacts tend to be more pronounced for high throughput data acquisition, which is typically desired in challenging projects^16^. Recent advances in image denoising and deconvolution can significantly mitigate these artifacts^21^.

The field of image restoration is a popular field in general image processing: state-of-the-art methods are based on multiresolution shrinkage^22, 23^, non-local pixel averaging^23, 24^, Bayesian estimation^25, 26^ or convolutional neural networks^27^. Even though most of these methods are available to the community, they are often impractical due to parameter finetuning and high computational demands. Similar to (semi-)supervised segmentation, users tend to be subjective and biased towards interactive image restoration in EM, suggesting that a ‘human-in-the-loop’ (HITL) can determine the optimal parameter settings for efficiency and reproducibility. Existing interactive frameworks^28–30^ tend to rely on parameters that are difficult to interpret and/or are computationally too intensive for efficient tuning and scaling towards large-scale 3D datasets. Furthermore, the current state-of-the-art in image restoration is evolving fast, prompting the need for a framework that is easily extendible with new algorithms. In this work, we propose a novel HITL framework called *DenoisEM* equipped with state-of-the-art image restoration algorithms combined with intuitive parameter interpretation available through ImageJ^31^ and computationally accelerated via a GPU-based back-end and high-level programming language called Quasar^32^. Our plugin is publicly available at http://bioimagingcore.be/DenoisEM.

## Results

### A human-in-the-loop (HITL) approach for semi-automated image restoration

Many solutions have been proposed for restoration of EM images^21^. However, practical solutions that allow for user feedback to apply state-of-the-art denoising on large 3D EM datasets generated by *e.g.* SBF-SEM or FIB-SEM are not readily available. Optimal finetuning of parameters in denoising is crucial w.r.t. the restoration result and this requires expert intervention. Automated parameter optimization approaches require a quality criterion (w.r.t. the task) and groundtruth, which is rarely available. We developed *DenoisEM*, a plugin with a HITL workflow that allows for efficient interaction and feedback by the expert. *DenoisEM* is a plugin for ImageJ^31^, an open source program that is extensively used in the microscopy community. The plugin allows for quick testing, comparison and application of different denoising solutions. The HITL plugin workflow (see figure 1) consists of six steps: data loading, initialization, region of interest (ROI) selection, noise estimation, interactive parameter optimization and final batch processing. Each step is automated as much as possible and user interaction is only required in the selection of the ROI and parameter settings.

**Figure 1.**
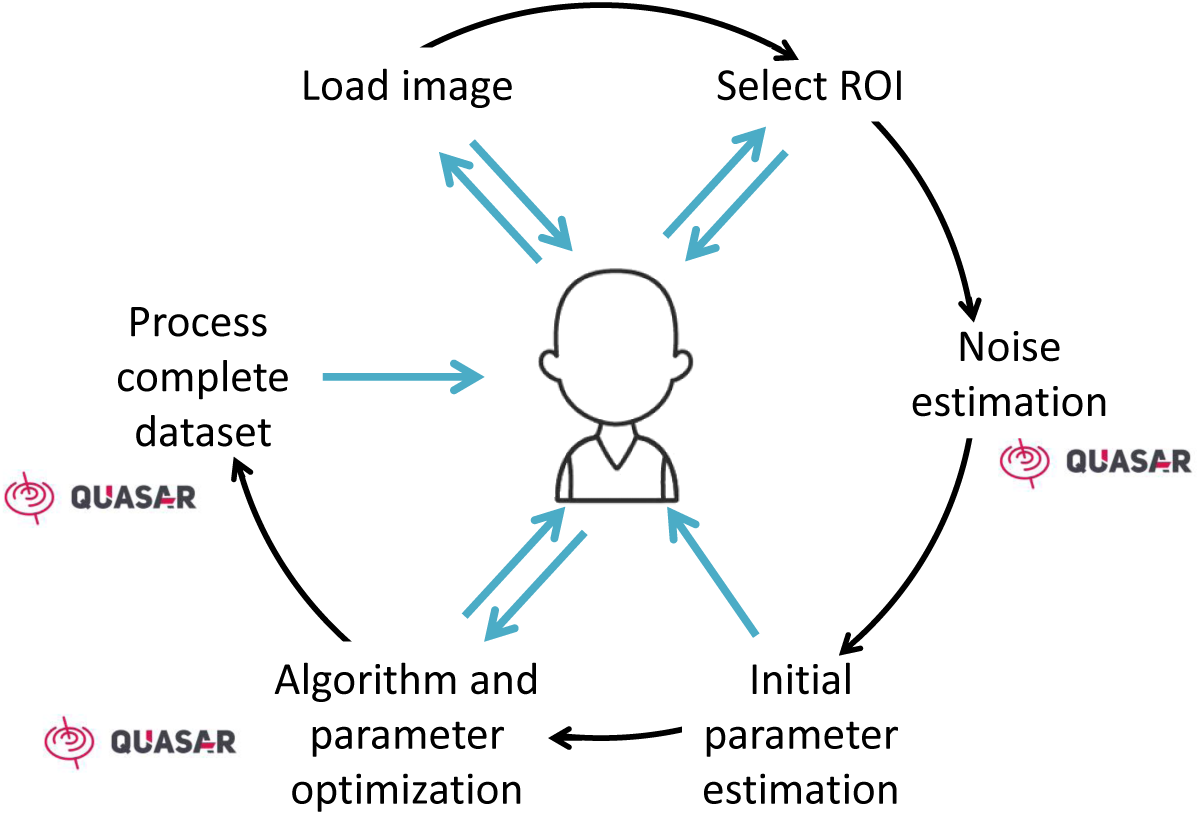
Graphical workflow of our proposed framework: an image is loaded and the computation back-end is prepared. Next, the user selects a ROI that is representative for the complete data set. The noise level is automatically estimated to derive near optimal parameter initialization. Next, the biological expert can optimize the parameter settings at a low latency visualization of the results according to his preferences (typically w.r.t. visualization and/or subsequent segmentation of specific objects). Once the optimal parameters for a specific algorithm are found, the complete data set is ready to be processed. The computationally intensive parts of the workflow are GPU accelerated and indicated with the Quasar logo^32^.

### DenoisEM provides an intuitive and interactive user interface

DenoisEM guides the user through the denoising process via a simple interactive user interface (see figure 2) in a step-by-step procedure:

- The user starts by opening an image (2D) or image stack (3D) in ImageJ. During startup of the plugin, this image is assigned as the reference image for restoration and all computational resources (CPU and, if available, GPU) are prepared (figure 2a).
- Next, the user can select a particular region of interest (ROI) in the reference image that is representative for the dataset and the analysis problem that follows restoration. This can be intuitively performed using the available selection tools in ImageJ. At this point, the noise level is automatically estimated on the ROI so that the initial parameter settings of the available algorithms are close to optimal and the required time for parameter finetuning can be minimized.
- The next step involves algorithm selection and interactive finetuning of the parameter settings. EM image data can be highly variable due to the used sample preparation technique, acquisition circumstances, cell type, *etc.* High quality image restoration therefore requires careful parameter tuning and algorithm selection w.r.t. the data. Each denoising algorithm has its advantages and disadvantages depending on the data and parameter settings can significantly influence the result (*e.g.* see supplementary Figures 1-9). Therefore, this is a crucial step that requires expert feedback. A strong asset of DenoisEM is that different algorithms (listed and briefly discussed in the Methods section), including recently developed ones, can be found in a single tool and are practical to use due to GPU acceleration. Switching between algorithms is done by checking the corresponding box and different parameters can be set by sliders. Typically the influence of different parameter settings is demonstrated at low latency, which facilitates the finetuning process. When visualization would lag, it is indicated by a red frame around the denoised image. To improve the efficiency of parameter optimization, tooltips indicate the influence of the parameter on the result (*e.g.* noise reduction or sharpening). When the user proceeds to a different algorithm or to the next step, the previous parameters are cached, making it feasible to switch back if necessary.
- In the final step, the user has the choice to apply the desired denoising algorithm and its corresponding parameters on the full image slice or stack. A new file is generated and the denoising algorithm and parameters are stored as metadata. These settings can then be applied to other datasets that require the exact same processing.

**Figure 2.**
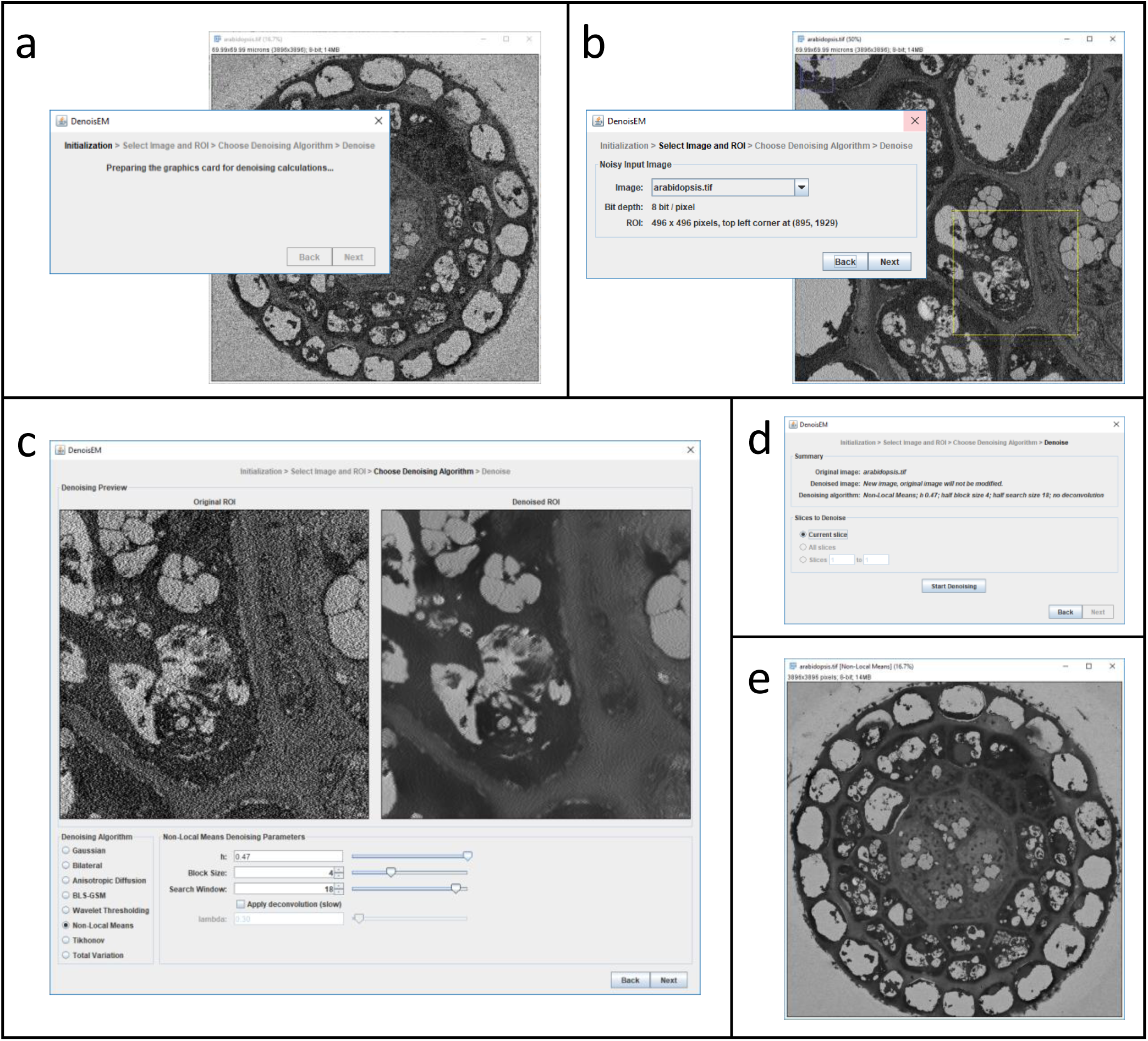
An overview of the user interface (UI) of the DenoisEM plugin for ImageJ. It is structured as a so-called ‘wizard’ that guides the user through the denoising process in a sequence of steps. (a) In ImageJ the user loads an image or image stack for denoising and starts the DenoisEM plugin. The UI wizard appears and the computational back-end for parallel processing on the CPU/GPU (Quasar) is initialized. (b) In the next step the user chooses any of the open images for denoising and selects a ROI on which denoising will be previewed. (c) Next the main panel in the plugin appears. At the top it shows side-by-side the noisy original as well as the denoised version of the selected ROI. In the bottom left corner the user can select one of eight denoising algorithms. The bottom right has controls for specifying the algorithm parameters. Typically, if the algorithm or its parameters are changed, the denoised ROI at the top is updated virtually instantaneously. This allows the user to easily assess the effect of algorithms and parameter settings. (d) After optimal settings are chosen, the user is shown a short summary and (for image stacks) can select the image slices that need to be denoised. During denoising the user is shown progress feedback. (e) When denoising is finished, a new image or image stack is created and displayed. The original image (stack) is left untouched.

We provided eight different denoising algorithms in DenoisEM. Supplementary figures 1-9 show the effect of applying each of these denoising algorithms under several different parameters on the same noisy image patch. The plugin user manual, available through Supplementary Note 2, describes the different algorithms and the associated parameters from a user perspective.

Because the DenoisEM plugin allows to select any image that is open in the program, it allows for alternating with other applications in ImageJ. After each step in the workflow, it is possible to go back to previous steps and at no point the original image is overwritten, to assure that the original image is never lost.

### DenoisEM improves visualization of 3D EM ultrastructure

We used DenoisEM on SBF-SEM images of an *Arabidopsis thaliana* root tip. The *en bloc* stained sample was prepared as described by Fendrich et al.^33^ and individual images of the 3D stack were acquired. The original image contained a significant amount of noise and we applied Tikhonov deconvolution (*λ* = 1.5, *σ* = 0.31 and *N* = 86 iterations) with the DenoisEM plugin. Visual assessment of the denoising effect by a biological expert was crucial to assure that no artifacts were introduced. Figure 3 shows the result of the denoising on four different ROIs from the datasets, and for each ROI both an XY and a YZ orthogonal view is shown. In this particular case, mild denoising was applied to avoid the loss of structural details in the image. Nevertheless, there was an effect on image quality and improved recognition of certain subcellular structures, *e.g.* the nuclear membrane (Figure 3 ROI 1) or endoplasmic reticulum (Figure 3b, ROI 3). Note that the image quality has also improved in the axial direction, even though the plugin performs lateral slice-by-slice denoising. We found that the noise level was decreased by two orders of magnitude (Figure 3c) using a state-of-the-art noise level estimator^34^.

**Figure 3.**
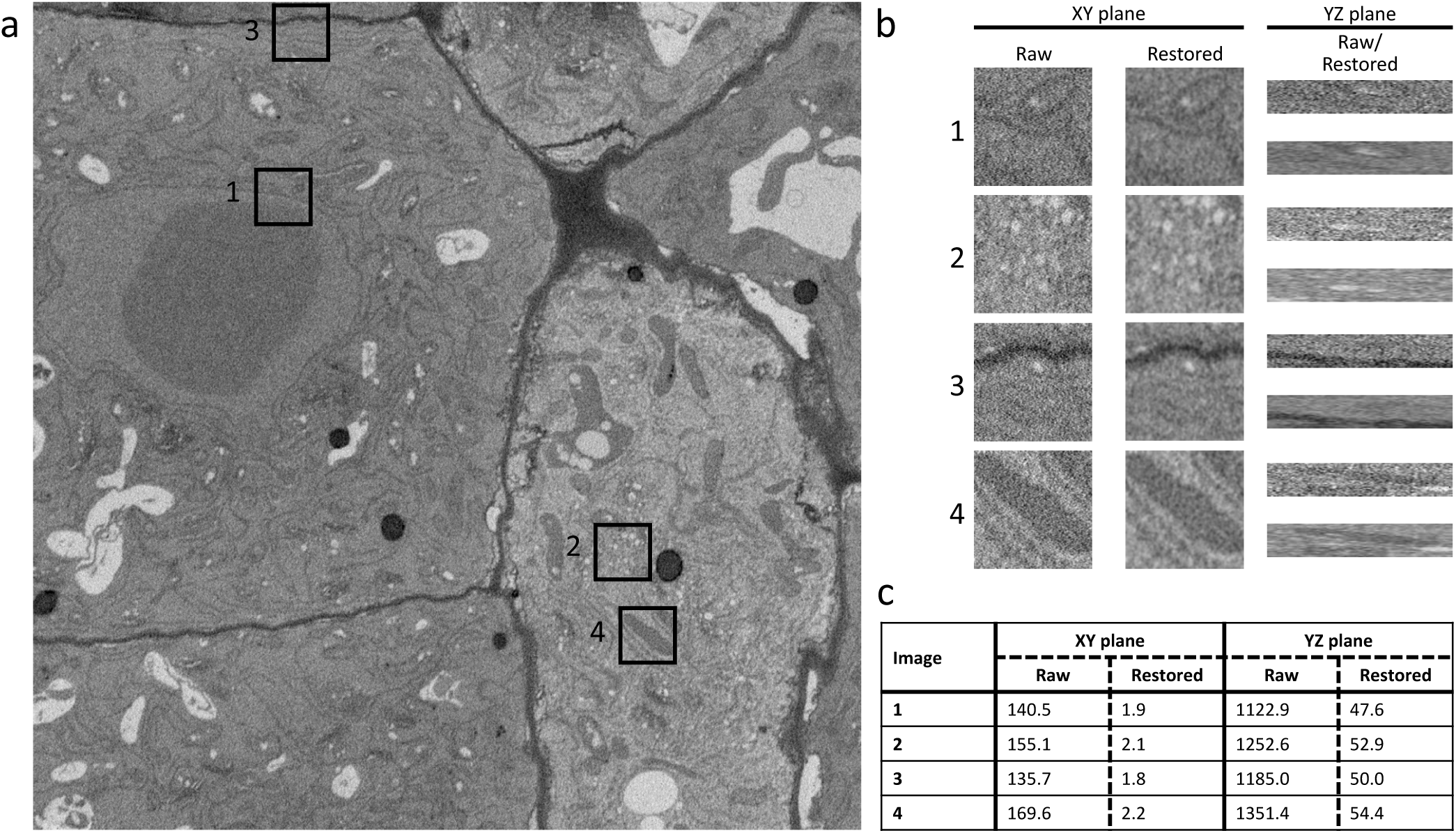
(a) Single XY section from an *Arabidopsis thaliana* root tip SBF-SEM acquisition and (b) four cross-sections of 150 × 150 × 27 ROIs that show the restored result from DenoisEM using Tikhonov deconvolution. For each ROI we show an XY and YZ section to illustrate that the image quality also improves along the *Z* direction, even though DenoisEM restores each XY slice independently. (c) For each ROI, we provide an estimation of the noise standard deviation 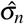^34^ in the raw and denoised patch to illustrate the image quality quantitatively. Note that the noise level decreases by almost two orders of magnitude.

In a next example we used SBF-SEM images of mouse heart tissue, prepared and imaged as described in Vanslembrouck *et al*.^35^ Figure 4a shows an unprocessed 2D image of heart smooth muscle cells. The sarcomeres can be observed with the A-bands as darker zones and the I-bands as brighter zones. We applied the anisotropic diffusion algorithm (step size *η* = 0.07, *N* = 6 iterations and a diffusion factor *κ* = 0.18) on this data using DenoisEM. The effect of denoising is shown in figure 4b, while figures 4e and f show the corresponding intensity histogram before and after denoising. Denoising is beneficial for the interpretation of the sarcomere organisation, since the A- and I-bands can be better distinguished after noise removal. This is illustrated in figure 4c and d, where intensity thresholding is used to generate a red and green mask for the A- and I-bands, respectively.

**Figure 4.**
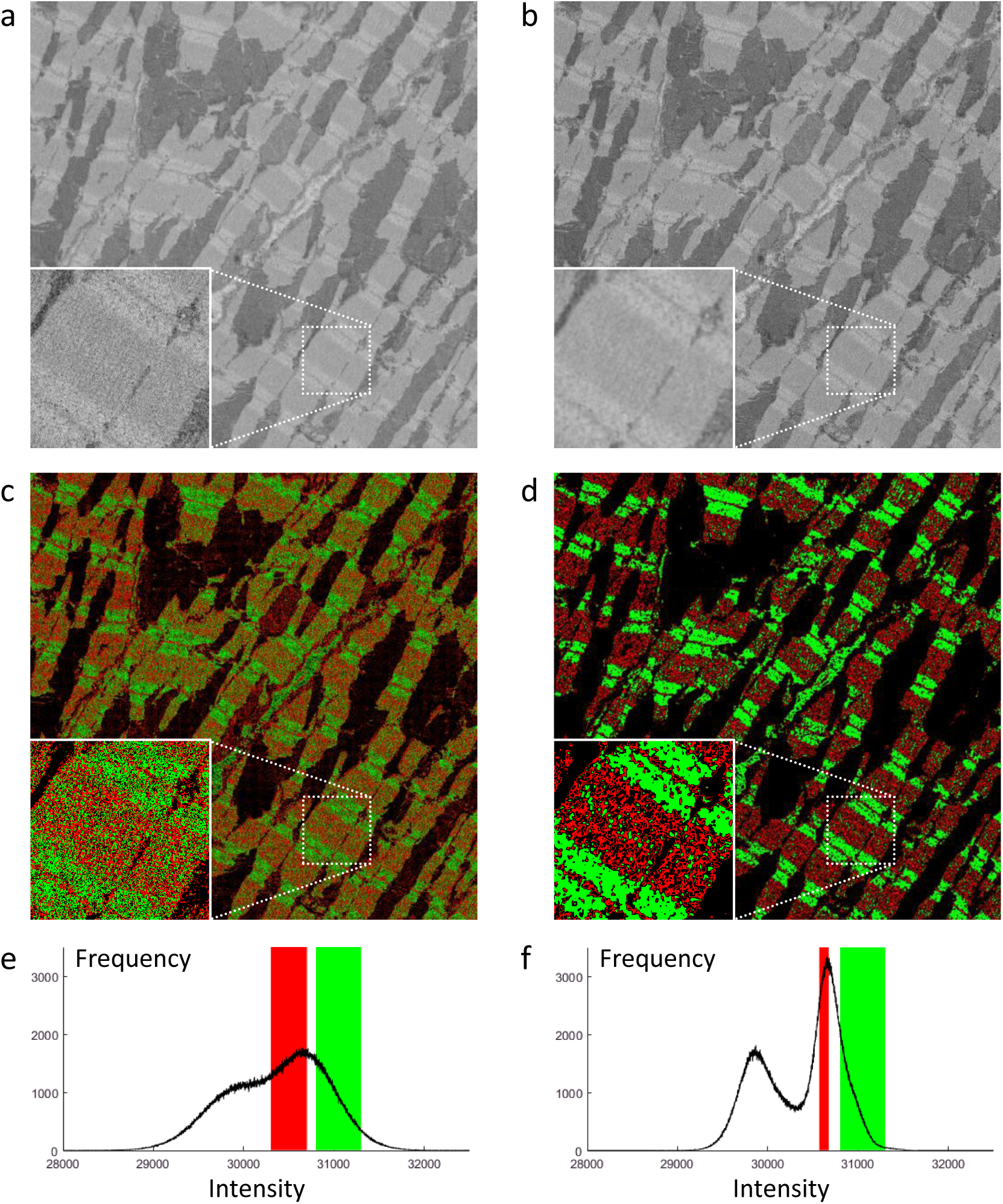
Mouse heart tissue was prepared for SBF-SEM. (a) One image of an SBF-SEM datastack is shown. The inset shows a zoom on the region indicated by the box. (b) The same image is shown after applying denoising. For visualisation, thresholding was applied for 2 ranges of intensity values and a red mask was used to indicate A-bands, a green mask to indicate I-bands. (c) and (d) show the masks for the raw and denoised data, respectively. (e) and (f) show the intensity histograms that correspond to images (a) and (b), with indication in red and green of the threshold values that were used to generate images (c) and (d).

### DenoisEM improves segmentation quality and speeds up image analysis

Mouse heart tissue was prepared as in Vanslembrouck et al.^35^ and imaged using FIB-SEM. In this particular dataset the lateral view shows a transverse filament section (figure 5a). The noise in the image is removed with DenoisEM by applying the non-local means algorithm with damping parameter *h* = 0.23, half window size *B* = 6 and half search window size *W* = 9 (figure 5d). For both the raw and denoised data, a rendering mask was created by intensity-based thresholding (figures 5b and e) with the ImageJ 3D Viewer (figure 5c and f). Denoising is crucial for the 3D visualisation of the filaments as separate objects. This is also demonstrated when performing object counting to determine the amount of individual filaments in an image. Figure 5g and i show the indicated regions from figure 5a and d, the raw and denoised XY view, respectively. Two experts counted the number of filaments in these ROIs three times, using the ImageJ ‘Cell Counter’ plugin. Each object is identified with a click and indicated with a yellow point. In the raw image, the filaments are less obvious to recognize and the number of indicated objects was lower as compared to the denoised image (figure 5k). When using a thresholded image to perform automatic segmentation and counting with the ImageJ ‘Analyze Particles’ tool, the number of counted objects in the denoised image corresponds to the manual counting, especially when applying a watershed filter prior to ‘Analyze Particles’ (figure 5j and k). The same analysis was performed on the raw image, resulting in values that are off by a factor of more than 1.7 compared to manual counting (figures 5h and k). This demonstrates that the raw images are not suited for automatic object counting and that the introduction of expert-guided denoising can prepare the images for an automatic analysis workflow. Here, the analysis by thresholding, watershed and the ‘Analyze Particles’ tool required 20 seconds per image on average, while manual counting required at least 2.5 minutes.

**Figure 5.**
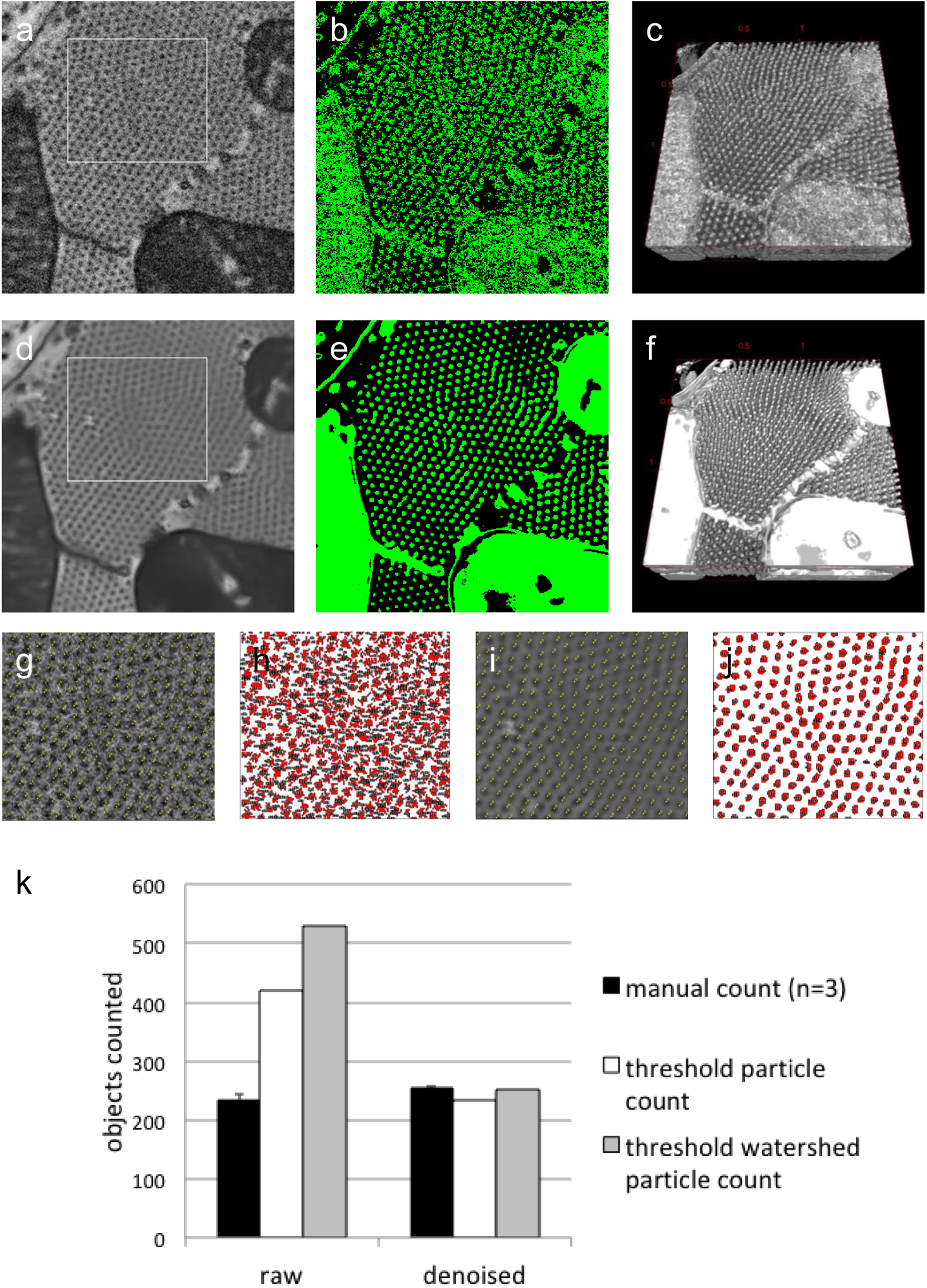
A 3D ROI from a FIB-SEM dataset, acquired at isotropic 5 nm resolution, from mouse heart tissue was used for denoising. (a) An XY view of 330 × 300 pixels of the raw image. (b) The image was used for creating a mask (in green) by intensity thresholding. (c) This mask was used for 3D rendering with the ImageJ 3D Viewer. (d) The data was denoised with DenoisEM, using the NLM algorithm and (e)-(f) segmented by thresholding. (g) The ROI indicated in panel (a) was used for manual counting of sarcomeres with the ImageJ Cell Counter Plugin. (h) Each item counted is indicated with a red dot. Panels (i) and (j) shows the counting results on the denoised data. (k) A graph showing the number of sarcomeres counted, either manual by three different individuals or by the ImageJ Analyse Particles tool on a thresholded image, with or without watershed.

To illustrate how our plugin can improve automated segmentation quality, we have performed experiments with the CREMI challenge dataset (https://cremi.org/) where volumes of serial section EM of the adult fly brain are considered for neuron reconstruction. Specifically, we use the pixel classifier from the ilastik framework19 to predict neuronal membranes. The prediction uses intensity, edge and texture features and a random forest pixel classifier combined with a small number of annotated pixels: approximately 4, 000 and 7, 000 membrane and background pixels, respectively (see top-left image in figure 6). To evaluate the influence of noise and denoising, we simulate noise of variable levels (*i.e. σn* = 0.01, 0.05, 0.1, 0.2) on the original (approximately noise-free) data and denoised these noisy datasets with the non-local means algorithm with a half window size *B* = 4, half search window size *W* = 5 and damping parameter *h* = 0.05, 0.15, 0.4, 0.7, respectively. Expert intervention was necessary to finetune the damping parameter w.r.t. neuronal membrane preservation for each noise level. Segmentation is performed on both the noisy and denoised datasets using the same annotated pixels. Figure 6 shows the generated probability maps and segmentation masks and figure 7 illustrates the quantitative results in relation to the noise level by means of the Dice coefficient (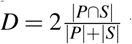 where *P* and *S* are the set of predicted and true foreground/background pixels, respectively). The results show that even a more advanced segmentation algorithm such as random forests pixel classification is not robust enough, as segmentation quality increases whenever denoising is performed as a pre-processing step. This is especially notable for higher noise levels, which is often the case when a short acquisition time was used (*e.g.* due to time or sample restrictions). Given that the pixel dwell-time is inversely related to the noise level *σ* in the image, it can be inferred from figure 7 that equal segmentation performance is achievable by accelerating the image acquisition by a factor of 20 (*σ* = 0.01 vs. *σ* = 0.2) and including denoising. The segmentation results even improved for a 10-fold acquisition acceleration (*σ* = 0.01 vs. *σ* = 0.1) when denoising is included. Practically speaking, this implies that acquisition times can be reduced from hours to minutes without sacrificing segmentation quality. Note that the denoising itself does not require a significant amount of overhead: *e.g.* the processing time for non-local means on a single CREMI dataset volume (1250 × 1250 × 125 pixels) requires less than a minute with a modern GPU.

**Figure 6.**
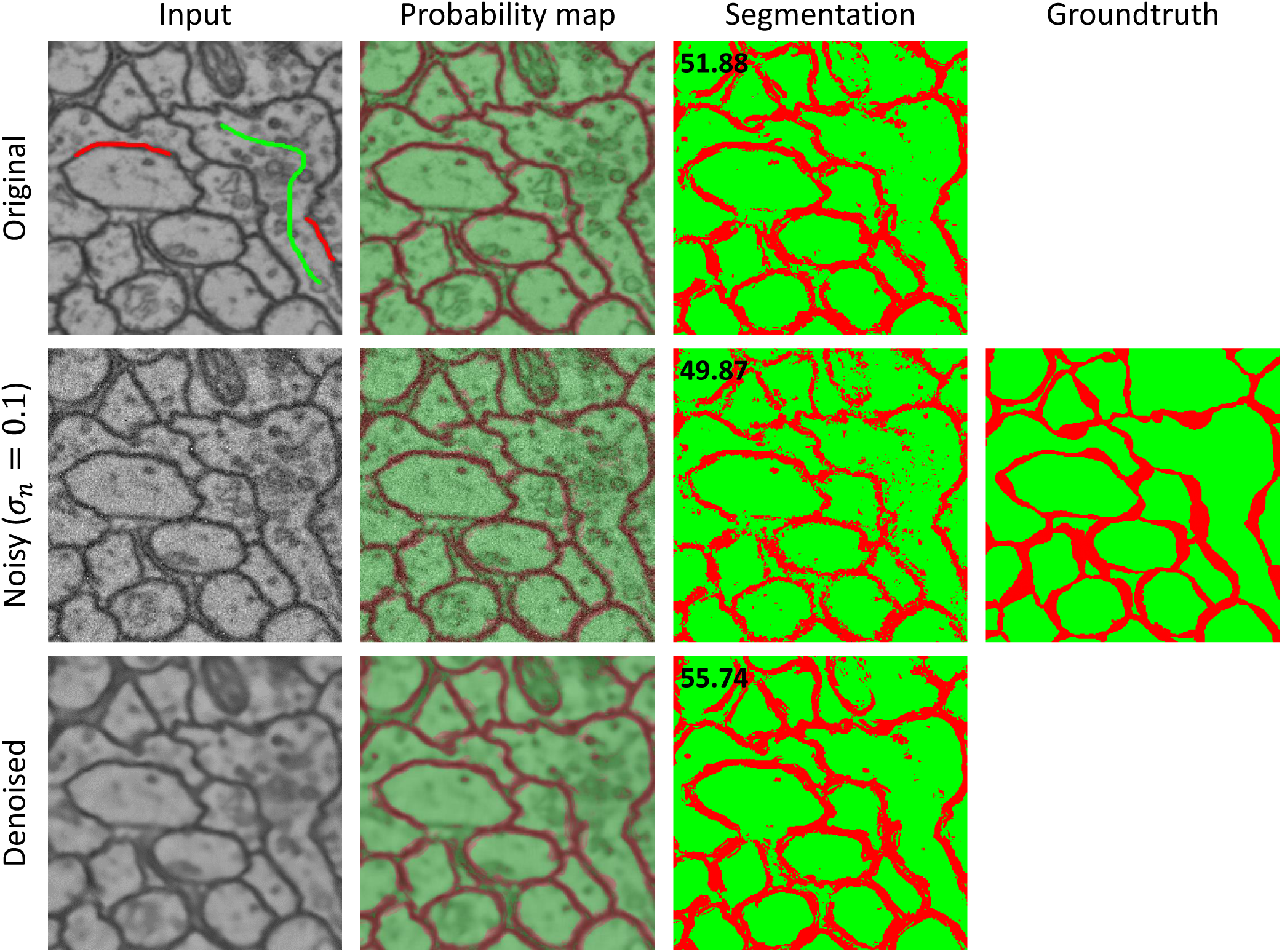
Segmentation results of automated neuron membrane segmentation using ilastik^19^ on the original CREMI data, with simulated noise (*σ*_*n*_ = 0.1) and the corresponding denoised version (non-local means with a half window size *B* = 4, half search window size *W* = 5 and damping parameter *h* = 0.4). The first column shows the input, the top image additionally shows a fraction of the labels that were used for training. The second column shows the probability output map of the random forest pixel classifier from ilastik. The third column shows the segmentation result by thresholding the probability maps and the fourth column the groundtruth segmentation (the Dice coefficient is shown in the upper left corner). Noise artifacts are clearly visible in the segmentation and can be avoided by denoising as a pre-processing step. Notice that denoising can even improve the segmentation result.

**Figure 7.**
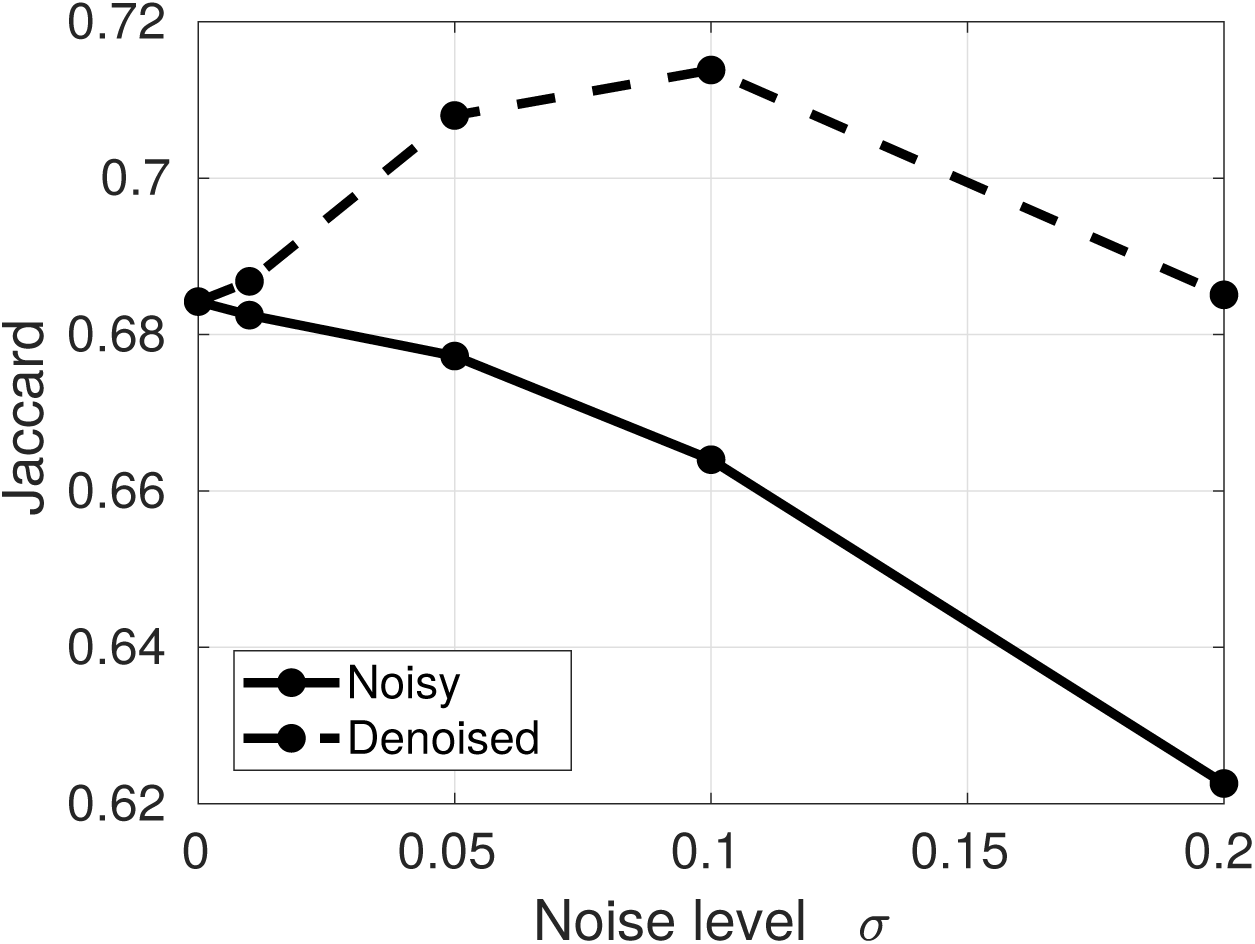
Segmentation quality (Dice coefficient) of the automated neuron membrane segmentation in the CREMI dataset using ilastik^19^. As the noise level in the data increases, the segmentation quality is significantly influenced whereas denoising pre-processing stabilizes (and even improves) the segmentation quality.

### DenoisEM performs at low latency and is easily extensible with new methods

In order to allow for a user in the loop, it is necessary to compute restoration results at a low latency, so that the user can test different parameter settings as fast as possible. The computational back-end of DenoisEM relies on a freely available programming framework called Quasar^32^, which reduces the complexity of heterogeneous programming on CPUs and GPUs to the programming in a high-level language like MATLAB/Python without significant runtime influence. Quasar is therefore ideal for development of novel image processing algorithms as prototyping is accelerated by parallel computing on the GPU and/or CPU.

As the plugin’s host application (ImageJ) is implemented in the Java programming language, we developed a bridge between Java and Quasar to connect front and back-end. This bridge uses the Java Native Interface (JNI) to wrap Quasar objects and functions in Java equivalents, providing an abstraction layer that separates the complexity of the Quasar C++ API from the DenoisEM plugin. The bridge is DenoisEM-agnostic and by itself a useful general building block for other applications wishing to leverage Quasar from within a Java environment. The DenoisEM plugin and the source code for the plugin, the Java-Quasar bridge and the denoising algorithms are freely available for non-commercial use.

We have compared the Quasar algorithm implementations available in DenoisEM to existing open-source implementations, which are typically not GPU accelerated. Figure 8 shows a comparison for bilateral filtering^36^, anisotropic diffusion^37^, BLS-GSM^22^ and non-local means denoising^24^ (results of the remaining methods can be found in the supplementary material). The results show that the Quasar implementations are one to two orders of magnitude faster than the classical CPU-based implementations. Considering a latency threshold of 100 ms (*i.e.* 10 FPS), this allows real-time processing of megapixel images for the bilateral filter, anisotropic diffusion and the more recently proposed non-local means algorithm. State-of-the-art denoising algorithms such as BLS-GSM used to take seconds to minutes for only a few megapixel sized images, whereas our Quasar implementation requires up to one second for 16 megapixel images. This allows for much faster parameter tuning by the expert. Note that this acceleration can also be obtained by incorporating *e.g.* CUDA libraries in the baseline implementations; however, this approach would require more development time and we believe that the high-level nature of Quasar is more scalable to this end. For example, expert CUDA developers required three months to implement an MRI algorithm^45^, whereas a single Quasar developer achieved the same numerical results at the same computational performance within a week. Additionally, we can observe that the obtained GPU speedups increase for larger inputs, which is desirable for large-scale computing. This is due to the fact that more pixels can be processed in parallel, and bounded by the amount of cores in the available GPU hardware.

**Figure 8.**
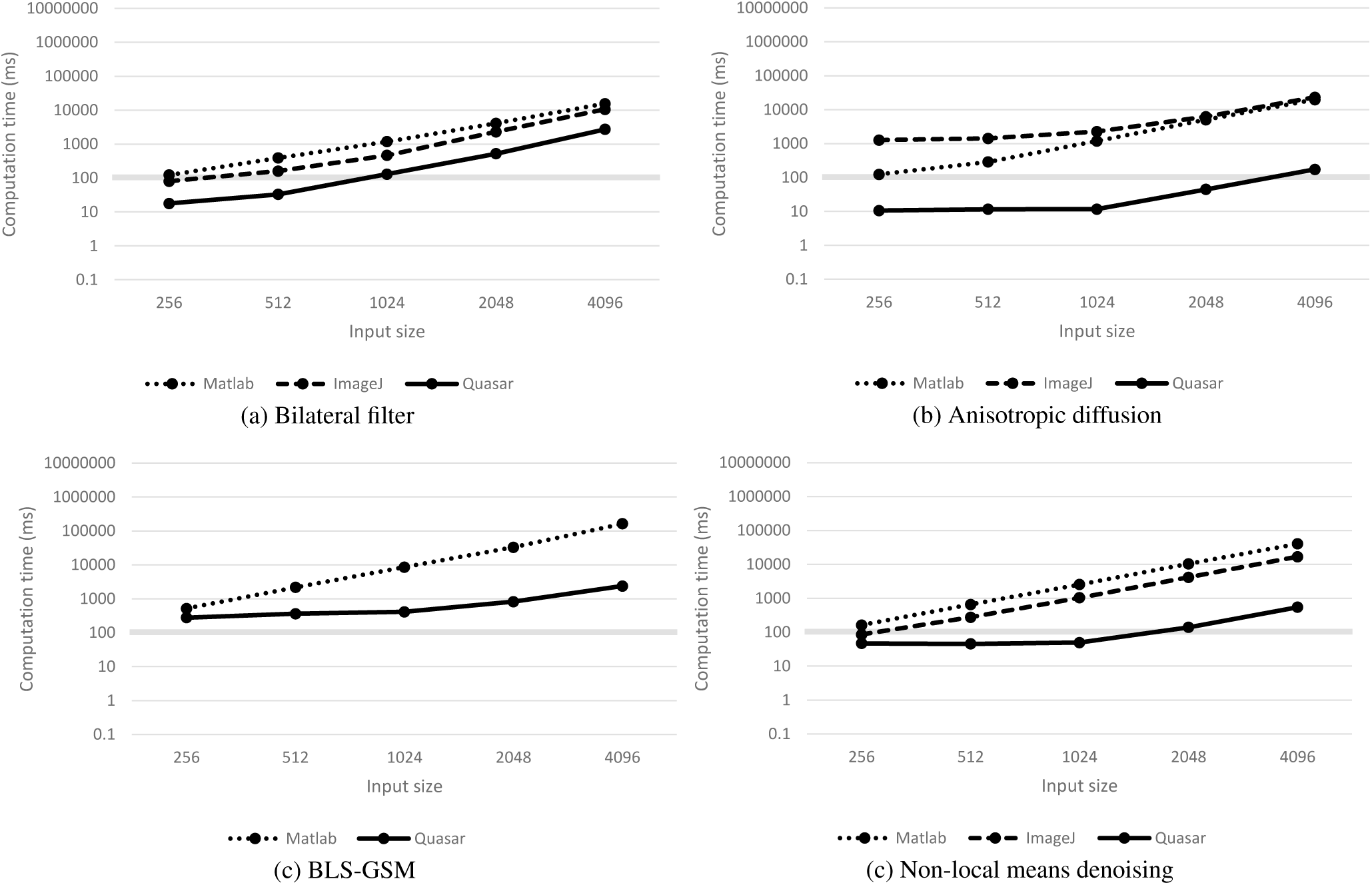
Absolute computational performance (in milliseconds) of bilateral filtering^36^, anisotropic diffusion^37^, BLS-GSM^22^ and non-local means denoising^24^ for different input sizes. A comparison is made between our proposed GPU-based Quasar framework and alternative implementations that are available to the scientific community: bilateral filter (ImageJ^38^ and MATLAB^39^ based), anisotropic diffusion (ImageJ^40^ and MATLAB^41^ based), BLS-GSM (MATLAB^42^) and non-local means denoising (ImageJ^43^ and MATLAB^44^ based). For each algorithm, we consider inputs of 256^2^, 512^2^, 1024^2^, 2048^2^ and 4096^2^ pixels. In general, our Quasar implementation performs 10 to 100 times faster compared to the existing software packages. Notice that the obtained speedup increases as the input image size increases due to the fact that GPUs are able to process more pixels in parallel.

## Discussion

In this work, we present DenoisEM, an ImageJ plugin that introduces the expert in the automatic workflow (HITL) by allowing for real-time parameter tuning. The HITL makes for a functional and efficient solution, but because the ultimate purpose can be diverse, there is no one-fits-all algorithm. The broad range of denoising possibilities offered here in one solution, make DenoisEM very versatile. The efficiency is due to the expert making fast judgements of the outcome of specific denoising parameters, the result being displayed at low latency on a ROI, thanks to GPU acceleration. Efficiency is also assured by allowing the user to go back and forth between the four different steps of the process. Reproducibility is also guaranteed by saving the algorithm parameter settings as metadata. This can also be useful in a later stage, *e.g.* for other datasets that require the same processing.

We validated the potential improvements that DenoisEM can provide in 3D EM image interpretation by denoising SBF-SEM image data of an *Arabidopsis thaliana* root tip. Experts confirmed that structures such as the nuclear membrane and endoplasmic reticulum were easier to recognize in the denoised data by optimally tuning the denoising parameters. As a second use-case, we used DenoisEM on noisy SBF-SEM data of mouse heart tissue and improved the visualization of sarcomeres as the A and B-bands were better separated.

We assessed the effect of denoising on 3D EM data as a processing step prior to automated segmentation, both intensity-based thresholding and pixel-level higher-order feature classifiers. An interesting conclusion is that segmentation quality does not significantly decrease for denoised noisy inputs compared to those that are noise-free. Consequently, the acquisition time can be shortened to increase throughput or avoid overexposure, without significantly affecting the subsequent segmentation.

We believe that DenoisEM is a plugin, accessible to the life science community, that can significantly improve the quality of large-scale 3D EM datasets and increase the throughput of EM imaging experiments. Although we focus our plugin on EM stacks, it can also be used for any (2D/3D) grayscale image producing modality (*e.g.* medical or astronomical data). Future work in HITL image restoration will be focused on predictive parameter optimization based on supervised regression models and analyzing correlations between the parameters of different algorithms. By keeping track of the EM metadata (*e.g.* modality, cell type, acquisition time, *etc.*) and the eventual objective (segmentation or visualization), we believe that more optimal parameters can be estimated, further minimizing manual intervention. Additionally, we will also extend the DenoisEM framework to multichannel image data, to include the light microscopy community. Moreover, the high-level nature of Quasar allows us to easily implement novel restoration algorithms (*e.g.* based on convolutional neural networks) that could further improve the restoration quality.

## Methods

### Sample preparation

Mice were maintained in standard specific pathogen-free (SPF) housing according to the European rules on animal welfare at the animal facility of the Center for Inflammation Research at VIB (Ghent, Belgium). The study was approved by a local ethics review board. Mice were anesthetized by intraperitoneal injection of ketamine/xylazine (70mg of ketamine and 10mg of xylazine per kg of body weight) and perfused, first with PBS containing heparin (20units/ml) for 2 minutes, followed by 2% paraformaldehyde (PFA; AppliChem, Darmstadt, Germany), 2.5% glutaraldehyde [Electron Microscopy Sciences (EMS), Hatfield, PA USA] in 0.15M cacodylate buffer (Sigma-Aldrich, Overijse, Belgium), pH 7.4, for 10 minutes. Next, heart muscle tissue was isolated and fixed overnight using 2% PFA, 2.5% glutaraldehyde in 0.15M cacodylate buffer, pH 7.4. Samples were thoroughly washed in 0.15M cacodylate buffer, pH 7.4, before small blocks were dissected to proceed with the staining protocol. Post-fixation was performed by incubating tissue-blocks in 1% osmium (EMS), 1.5% potassium ferrocyanide (EMS) in 0.15M cacodylate buffer, pH 7.4.

For SBF-SEM, post-fixation with osmium was followed by incubation in 1% thiocarbohydrazide (TCH; EMS) and subsequent washes in double-deionized water (ddH2O). Next, a second incubation in 2% osmium in ddH2O was performed. Both TCH and the second osmication were repeated after this. The samples were then washed in ddH2O and placed in 2% uranic acetate (UA; EMS). After the following washing step, Walton’s lead aspartate staining was performed for 30 minutes at 60°C. For this, a 30mM l-aspartic acid solution was used to freshly dissolve lead nitrate (20mM, pH 5.5) just before incubation.

For FIB-SEM, the fixed tissue-blocks were washed in ddH2O for four consecutive steps, refreshing the ddH2O after every step. Next, incubation in 1% osmium in ddH2O was followed by washing in ddH2O, incubation in 1% UA and again washing steps with ddH2O.

After the final washing steps, samples for both FIB-SEM and SBF-SEM were dehydrated using solutions of 50, 70, 90, and twice 100% ethanol. Samples were then placed in 100% acetone and embedded in Spurr’s resin (EMS) by incubation in 50% Spurr’s in acetone, followed by 4 incubations in 100% Spurr’s. Polymeration was done overnight at 60°C. Except for the Walton’s lead staining all steps were performed using a Pelco Biowave Pro Microwave Tissue Processor (Tedpella Inc, Redding, CA USA). More detailed protocols can be found in Vanslembrouck et al^35^.

For SBF-SEM, the sample was mounted onto an aluminum pin, trimmed into a pyramid shape using an ultramicrotome (Leica, Ultracut) and the block-surface was trimmed until smooth and at least a small part of tissue was present at the block-face. Next, samples were coated with 5 nm of platinum (Pt) in a Quorum Q 150T ES sputter coater (www.quorumtech.com). The aluminum pins were placed in the Gatan 3View2 in a Zeiss Merlin SEM. For FIB-SEM samples were mounted on Aluminum stubs (EMS) and coated with 10 nm of Pt.

### Image acquisition

For SBF-SEM acquisitions a Zeiss Merlin with Gatan 3View2 was used. Acquisition parameters for images of heart tissue were acceleration voltage of 1.7 kV and 100 pA, dwell time of 1 *µ*s. Images were collected at 8000 × 8000 pixels with a pixel size of 12.44 nm and slicing was done at z-steps of 50 nm. For image reconstructions a ROI of 1500 × 1500 pixels was chosen and its 101 consecutive images. Acquisition parameters for images of *Arabidopsis thaliana* root tips were collected at acceleration voltage of 1.6 kV and 80 pA, dwell time of 1 *µ*s. The pixel size was 13 nm and the slice thickness 70 nm. FIB-SEM imaging of murine heart tissue samples was done with a Zeiss Crossbeam 540 at 5 nm pixels and slicing at 5 nm sections.

### Implemented restoration methods

In this section, we will give a brief overview of the implemented algorithms. This includes the median absolute deviation (MAD) noise estimator, least squares parameter estimator, Gaussian filtering (GF), bilateral filtering^36^ (BF), anisotropic diffusion^37^ (AD), wavelet thresholding^46^ (WT), Bayesian least squares Gaussian scale mixtures^22^ (BLS-GSM) and non-local means denoising^24^ (NLM) and deconvolution^47^ (NLMD). These methods are complementary in terms of computation time and restoration performance: algorithms such as BLS-GSM and NLMD are computationally more intensive than, for example, GF and AD, but the former are more likely to outperform the latter. This trade off between computation time and image quality can be tuned by the expert. In the supplementary material, we discuss the influence of the implemented methods and their respective parameters on the restored image. For a more in-depth description of these methods, we refer to the respective references of the algorithms.

We will denote 3D images as vectors by stacking the pixels using slice-by-slice raster scanning ordering. More specifically, the acquired image **y** ∈ ℝ^*N*^ (where *N* is the number of voxels in the image) is assumed to be degraded by blur and additive noise:

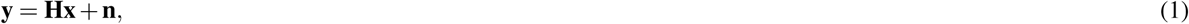

where **x** ∈ ℝ^*N*^ is the underlying, degradation-free image, **H** ∈ ℝ^*N*×*N*^ represents the microscope point-spread function (PSF) and **n** ∈ ℝ^*N*^ is a stochastic, noise term. The noise term is assumed to be mean-zero and with a constant variance, *i.e.* [**C**]_*i,i*_ = *σ* ^2^ (where **C** is the noise covariance matrix).

#### Noise estimation

Noise estimation involves the task of estimating the noise variance *σ* ^2^ based on the acquired image **y**. The method that is implemented in our plugin is the median absolute deviation (MAD) noise estimator:

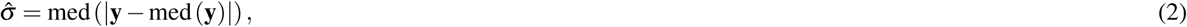

where med(·) denotes the median operator. The absolute difference of **y** and its median value provides a pixel-wise, noise-robust measure of how much the signal deviates from its expected value. Taking the median over these values therefore gives a robust estimation of the noise standard deviation over the complete image **y**.

#### Parameter estimation

Consider an algorithm **f**_***θ***_ (·) with parameters ***θ*** ∈ ℝ^*p*^ that maps a noisy image **y** onto a noise-free estimation 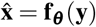. Based on the estimated noise level 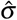 (see equation (2)), we estimate the optimal parameter settings through parameterized polynomial expressions w.r.t. a training dataset of noise-free benchmark EM images **x**_*k*_, for *k* = 1, *…, K* (*K* = 100 in our case). These images were degraded with Gaussian noise of different levels *σ*_*m*_ resulting in noisy images **y**_*k,m*_. The optimal parameters ***θ*** _*m*_ ∈ ℝ^*p*^ were determined for each noise level *σ*_*m*_ in least squares sense:

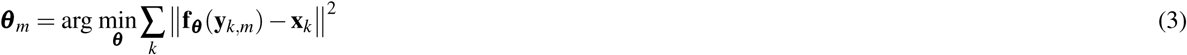

Next, the polynomial coefficient parameters **a**_*j*_ ∈ ℝ^*q*^ were optimized with a least squares fit:

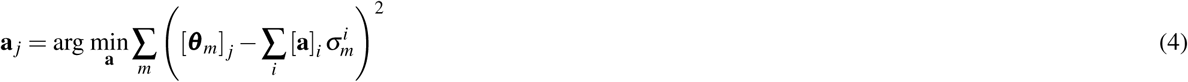

Finally, the estimated parameters that correspond to the estimated noise level 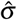 are computed as:

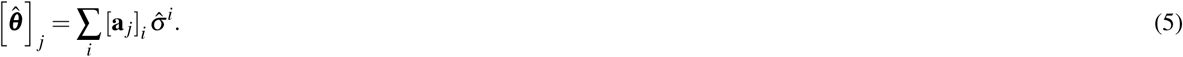

In practice, we concluded that a quadratic polynomial fit (*q* = 2) approximates the optimal parameters (in lease squares sense) sufficiently close.

#### Gaussian filter

The Gaussian filter is a special case of linear, smoothing filters which combine pixel values linearly in a local neighborhood in order to restore a pixel value. Practically, this comes down to a convolution of the noisy image **y** with a convolution kernel **G** (which is Gaussian in this case).

#### Wavelet thresholding

Wavelet transforms^48–51^ separate image information across multiple frequency scales and magnitudes. Noise tends to be spread among all the transformed coefficients whereas the coefficients that represent actual discontinuities typically stand out. Therefore, a popular method to reduce the amount of noise in an image is to reduce the magnitude of the transformed coefficients, this is also known as *wavelet shrinkage*^46^. More specifically, the restored image is found by transforming the image **y** to the wavelet domain, shrink the noisy coefficients and transform back to the spatial domain:

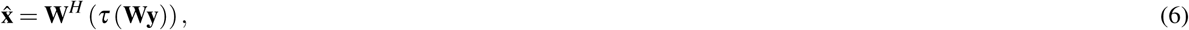

where **W** represents the used wavelet transform and **W**^*H*^ is its Hermitian transpose. The shrinkage function *τ* is typically the soft-thresholding operator:

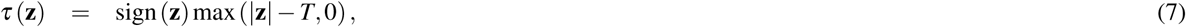

where all the functions operate component-wise and *T* is a thresholding parameter.

#### Anisotropic diffusion

The anisotropic diffusion filter^37^, commonly used in the context of EM image restoration^52, 53^, introduces non-linearity by describing linear filters in a partial differential equation (PDE) domain and extending it to a non-linear case. The true image **x** is embedded in a family of images **x**_*t*_, obtained by convolving the image **x** with Gaussian filters with a variance that increases with *t*. This diffusion process can be described by the so-called linear *diffusion heat equation*^54^:

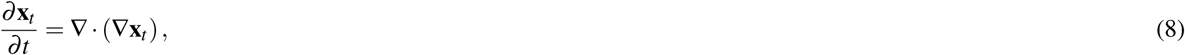

where ∇ represents the gradient operator with respect to the spatial coordinates. This isotropic diffusion method ignores edge information and consequently blurs edges. The anisotropic diffusion filter mitigates this by integrating a gradient magnitude weighting function:

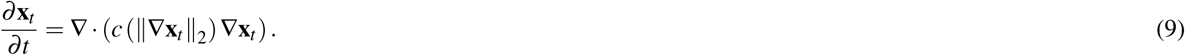

Two non-linear gradient regularization functions were proposed in Perona et al.^37^ and are currently still among the most commonly used:

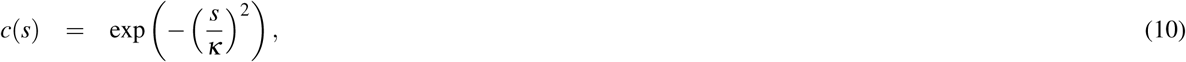

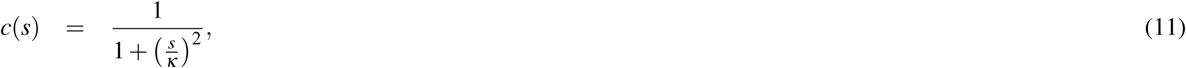

where *κ >* 0 is a parameter that offers a trade-off between noise reduction and edge preservation. The idea is that the function *c*(·) returns small values for large gradient magnitudes and vice versa such that edges are less diffused (*i.e.* blurred) and only in the direction of the edge (*e.g.* to avoid that horizontal edges will be blurred in the horizontal direction).

#### Bilateral filter

It is argued that local linear filters (such as the Gaussian filter) tend to oversmooth image edges and require non-linearities in order to obtain a better restoration estimate^55^. In former EM research^56, 57^ the bilateral filter^36^ is used: this is a locally adaptive spatial filter that avoids oversmoothing by averaging less aggressively along edges. More specifically, the restored *i*-th pixel 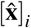 is computed as:

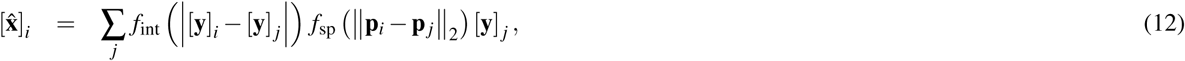

where *f*_int_, *f*_sp_: ℝ → ℝ are kernel functions that weigh the intensity and spatial distance, respectively, and **p**_*i*_ ∈ ℝ^3^ represents the 3D spatial position vector that corresponds to the index *i*. Similar to the Gaussian filter, pixels [**y**] _*j*_ nearby the reference pixel [**y**]_*i*_ will be assigned larger averaging weights through *f*_sp_. However, pixel intensities that differ very much from the reference pixel (typically edges) are assigned low weight values through *f*_int_, which leads to less blurring along edges.

#### Tikhonov deconvolution

Tikhonov restoration^58^ exploits the fact that images generally consist of many smooth regions separated by a much smaller amount of edges. As a consequence, the total edge magnitude in the restored image should be penalised, in other words:

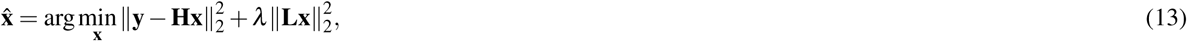

where **L** is typically a matrix that acts as a gradient or Laplacian operator in order to quantify edges in the image (we use the Laplacian).

#### Total variation deconvolution

The total variation prior (TV)^59^ assumes that natural images **x** should consist of flat regions delineated by a relatively small amount of edges. Mathematically, this is expressed as minimizing the total variation of the image:

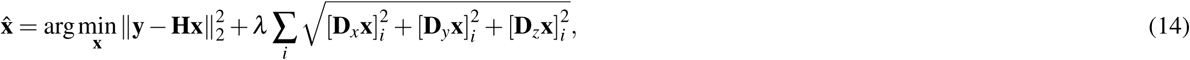

where **D**_*x*_, **D**_*y*_ and **D**_*z*_ are matrices that express the variation of the image along the *x, y* and *z* axis, respectively (*e.g.* first order derivative approximations).

#### BLS-GSM

The BLS-GSM method^22^ decomposes the image into *J* scales and *K* oriented pyramid subbands, denoises the highpass subbands and inverts the pyramid transform. An *M* × *M* neighborhood around a reference coefficient [**v**]_*c*_ of a subband is considered and represented as a vector **v**, by column stacking. These coefficients are then modeled as Gaussian Scale Mixtures (GSM), *i.e.* the product of a Gaussian distributed vector 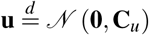 (where 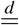 indicates distribution equality) and the square root of an independent, positive scalar random variable *z*:

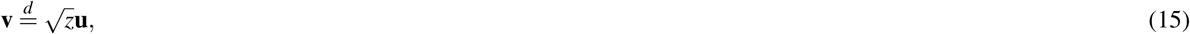

such that a noisy neighborhood patch **w** is described by:

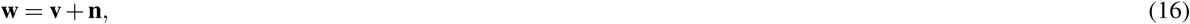

where 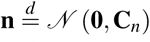 are the noise coefficients. Based on this model, the reference coefficient is approximated using the Bayesian least squares (BLS) estimator which reduces to^22^:

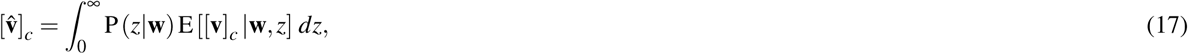

where P(*z*|**w**) is found with Bayes’ rule and E[**v**|**w**, *z*] is computed as a local linear Wiener estimate:

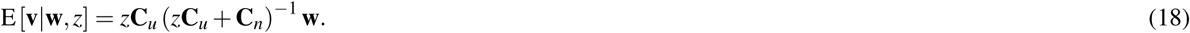

Mainly due to accurate statistical noise modelling, BLS-GSM has become the state-of-the-art in multiresolution-based denoising.

#### Non-local means denoising

Since the introduction of the non-local means (NLM) filter^24^, self-similarity denoising approaches gained a lot of interest because of their high performance. More specifically, the NLM algorithm estimates the noise free image pixels as weighted averages of acquired pixel values:

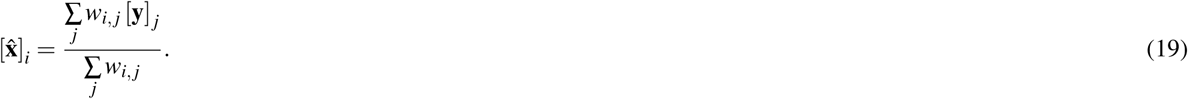

The self-similarity constraint turns up in the way the weights *w*_*i, j*_ are computed: these should be large (respectively, small) for similar (respectively, dissimilar) pixels *i* and *j*. Pixel similarity is defined along a local pixel neighborhood and searched for in both a local and non-local region. This is implemented in the originally proposed weights^24^:

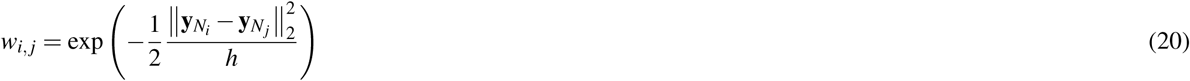

where 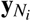 denotes a local neighborhood *N*_*i*_ of the acquired pixel [*y*]_*i*_ and *h* is a similarity damping parameter. In our plugin, we employ improved weights, proposed by Goossens et al.^60^

It has been shown that the NLM algorithm can be equivalently expressed by means of a Bayesian estimator with non-local image prior^26^. The work of^47^ proposes an NLM deconvolution algorithm by extending the Bayesian estimator with non-local prior of^26^ to a deconvolution estimator:

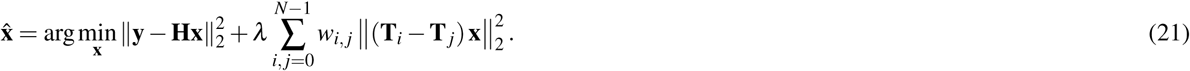

where *λ* is a regularization parameter, **T**_*i*_**x** is a vector with [**x**]_*i*_ on the first position and **H** is the estimated PSF of the microscope.

### Image restoration settings for the experiments

Most image restoration methods have parameters that may affect computational performance: *e.g.* search windows, block sizes, number of iterations, *etc.* For computational comparison, we selected parameter settings that corresponded to those that were most frequently preferred by biological experts during our experiments. In particular, we only report the parameters that affect computation time:

- Gaussian filter: window size 7 × 7 (fixed in the plugin)
- Wavelet thresholding: 3D dual-tree complex wavelet transform^61^, 6 scales (first scale filters by Abdelnour Farras, other scale filters using design from Selesnick^62^)
- Anisotropic diffusion: *N* = 5 iterations
- Bilateral filter: window size 15 × 15 (fixed in the plugin)
- Tikhonov denoising: *N* = 10 iterations
- Total variation denoising: *N* = 100 iterations
- BLS-GSM: *J* = 3 scales, window size 3 × 3 (fixed in the plugin)
- Non-local means denoising: half window size *B* = 4, half search window size *W* = 5
- Non-local means deconvolution: half window size *B* = 4, half search window size *W* = 5, *N* = 20 iterations

The algorithms were implemented in Quasar and compared to open source MATLAB-based (wavelet thresholding^63^, anisotropic diffusion^41^, bilateral filter^39^, Tikhonov denoising^64^, total variation denoising^65^, BLS-GSM^42^, non-local means denoising^44^, non-local means deconvolution^66^) and ImageJ-based (anisotropic diffusion^40^, bilateral filter^38^, non-local means^43^) implementations if available. The experiments were performed with an Intel(R) Core(TM) i7-4930K CPU @ 3.40GHz and an NVIDIA GeForce GTX 1070 GPU and repeated for various sizes of 2D input images (256^2^, 512^2^ up to 4096^2^).

SBF-SEM images, acquired with the Zeiss Merlin and Gatan 3View2 detector, are originally generated as 16-bit images. All image restoration techniques (denoising, registration, segmentation) were performed with floating point precision. The mouse lung artery data was restored using wavelet thresholding with a threshold value of *T* = 0.08. The Arabidopsis thaliana root tip was restored using Tikhonov deconvolution with the following parameters: *λ* = 1.5, *σ* = 0.31 and *N* = 86 iterations. The murine heart tissue dataset was processed with anisotropic diffusion with a step size *ν* = 0.07, *N* = 6 iterations and a diffusion factor *κ* = 0.18. ImageJ was used to apply thresholding on the raw and denoised image. The values of the selected intensities for the A-bands (shown in red in figure 4) was [30300, 30700] and [30589, 30674] for the raw and denoised image, respectively. The values of the selected intensities for the I-bands (shown in green in figure 4) was [30800, 31300] for both the raw and denoised image. The mouse heart tissue was denoised using non-local means with damping parameter *h* = 0.23, half window size *B* = 6 and half search window size *W* = 9. A reference area of 340×340 pixels and 117 sections was cropped and used in local template matching for registration. Reconstruction was performed similarly as with the murine heart tissue, using threshold values from the interval [32, 78]. As a final step a conversion to 8-bit was always done before exporting to PNG, to allow for final visualisation.

## Supporting information

All supplementary material

## Data availability

The DenoisEM plugin is available at https://bioimagingcore.be/DenoisEM. We stimulate the community to build on our work by open-sourcing the plugin. The raw data that was used for this manuscript is located at https://bioimagingcore.be/DenoisEM/data.

## Acknowledgements

This research has been made possible by the Agency for Flanders Innovation & Entrepreneurship (VLAIO), Grant Number: IWT.141703 and BOF, Grant Number: BOF15/PDO/003. We gratefully acknowledge the support of NVIDIA Corporation with the donation of the Titan X Pascal GPU used for this research. The Zeiss Merlin with 3View and Zeiss CrossBeam 540 were purchased by funding from the CLEM-grant and VIB Technology Fund.

## Author contributions statement

J.R., F.V., A.K., A. G. and S.L. conceived and designed the experiments. J.R., J.A., H.L. and B.G. implemented the denoising and registration algorithms. F.V. and B.G. developed the software plugin. All authors analyzed the results and wrote the paper.

