## Supplementary material for "A “Human-in-the-Loop” Approach for Semi-automated Image Restoration in Electron Microscopy": All supplementary material

### Supplementary material for the manuscript entitled: *A ‘‘Human-in-the-Loop’’ Approach for Semi-automated Image Restoration in Electron Microscopy*

This document provides supplementary material for the manuscript entitled: *A ‘‘Human-in-the-Loop’’ Approach for Semi-automated Image Restoration in Electron Microscopy*. In particular, we provide references to the raw data that was used for the experiments, a reference to our DenoisEM plugin, visual results of different parameter settings and computational performance metrics of all restoration methods in the plugin.

**Supplementary Note 1: data availability**

The data that was used to generate the results in the manuscript is available to the community on the locations specified below:

- Arabidopsis root tip (figure 3):
  - Raw: http://bioimagingcore.be/DenoisEM/data/arabidopsis-raw.tif
  - Restored: http://bioimagingcore.be/DenoisEM/data/arabidopsis-restored.tif
- Murine heart tissue (figure 4):
  - Raw: http://bioimagingcore.be/DenoisEM/data/murine-raw.tif
  - Restored: http://bioimagingcore.be/DenoisEM/data/murine-denoised.tif
- Mouse heart tissue (figure 5):
  - Raw: http://bioimagingcore.be/DenoisEM/data/mouse-raw.tif
  - Restored: http://bioimagingcore.be/DenoisEM/data/mouse-denoised.tif
- CREMI challenge data – fly brain (figure 6):
  - Full dataset: https://www.cremi.org
  - Subset: http://bioimagingcore.be/DenoisEM/data/flybrain/

**Supplementary Note 2: software**

The DenoisEM plugin can be downloaded at our project page http://bioimagingcore.be/DenoisEM. This location additionally provides software and hardware prerequisites, installation instructions, a getting started example and frequently asked questions. We have also provided a user manual, available at http://bioimagingcore.be/DenoisEM/doc/DenoisEM-manual.pdf, where the user can find practical information about the plugin for *e.g.* parameter finetuning. We are happy to help with any practical issues regarding the plugin and stimulate the community to contact us for this.

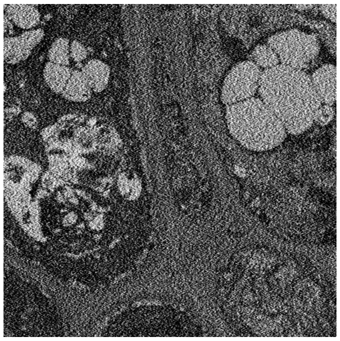

**(a)** Original noisy image.

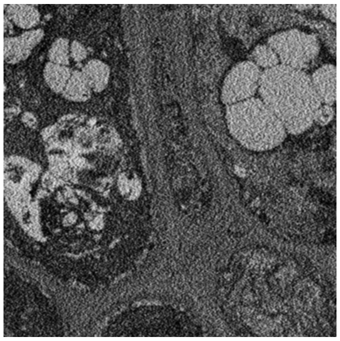

**(b)** Gaussian filter, $\sigma = 1.0$

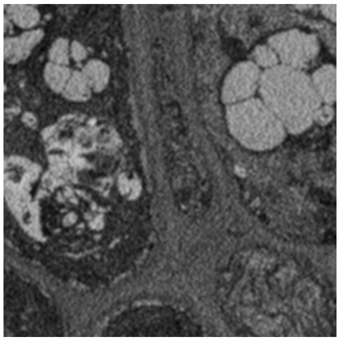

**(c)** Gaussian filter, $\sigma = 2.0$

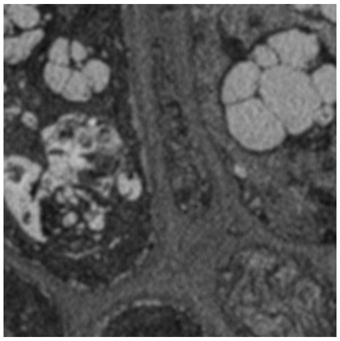

**(d)** Gaussian filter, $\sigma = 3.0$

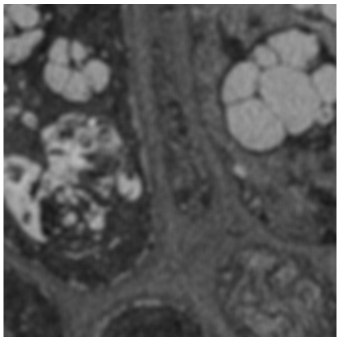

**(e)** Gaussian filter, $\sigma = 4.0$

**Supplementary Figure 1.** Denoising with Gaussian filtering. The original noisy image is shown on the left; on the right are the denoised results for increasing values of the blur kernel size $\sigma$. As $\sigma$ grows larger, the Gaussian filter reduces noise more aggressively at the risk of blurring edges.

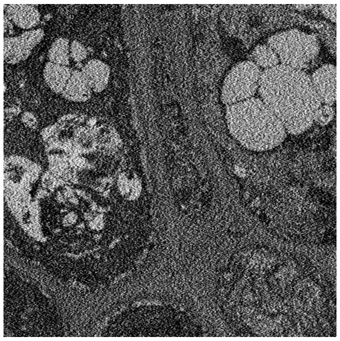

**(a)** Original noisy image

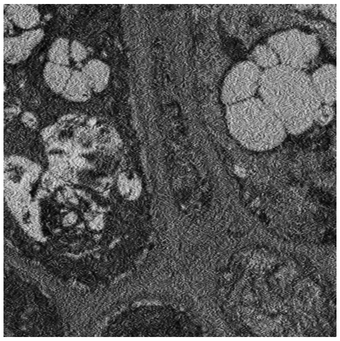

**(b)** Wavelet thresholding, $T = 0.25$

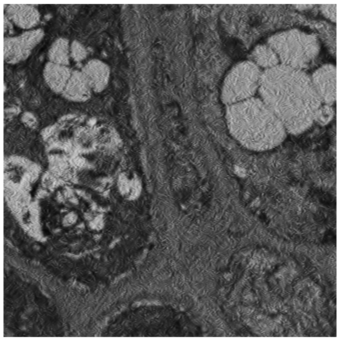

**(c)** Wavelet thresholding, $T = 0.5$

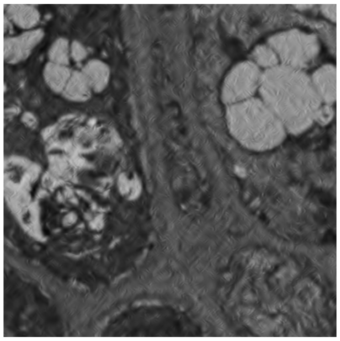

**(d)** Wavelet thresholding, $T = 1$

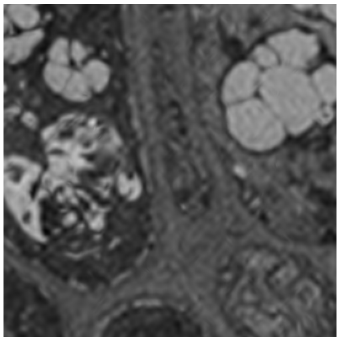

**(e)** Wavelet thresholding, $T = 2$

**Supplementary Figure 2.** Denoising via wavelet thresholding. The original noisy image is shown on the left; on the right are the denoised results for increasing values of the wavelet threshold parameter $T$. Increasing the thresholding parameter leads to higher noise suppression but also potential introduction of wavelet artifacts (*e.g.* ringing).

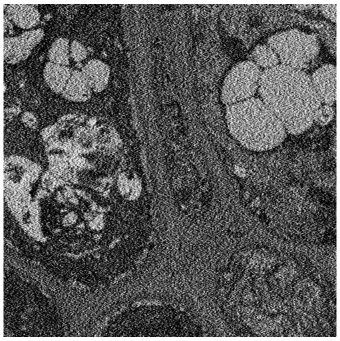

**(a)** Original noisy image

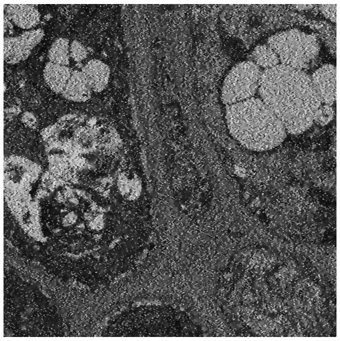

**(b)** $\kappa = 0.15$

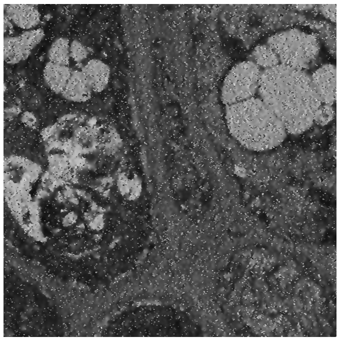

**(c)** $\kappa = 0.20$

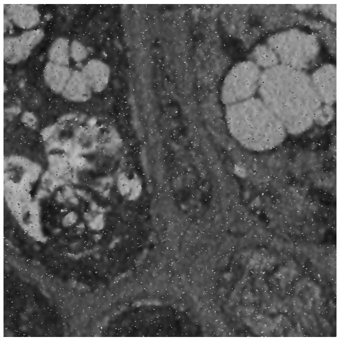

**(d)** $\kappa = 0.25$

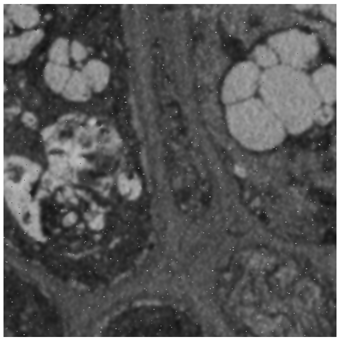

**(e)** $\kappa = 0.30$

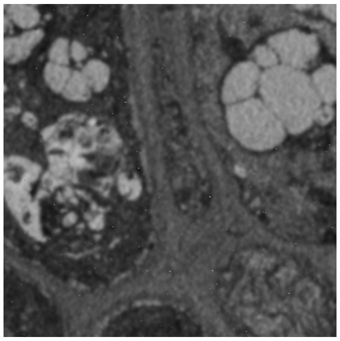

**(f)** $\kappa = 0.35$

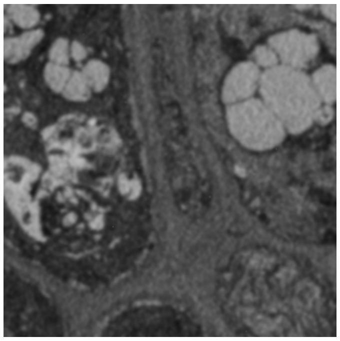

**(g)** $\kappa = 0.40$

**Supplementary Figure 3.** Anisotropic Diffusion denoising. The top image is the original noisy image; the rows below show the result of denoising with anisotropic diffusion for increasing values of the diffusion parameter $\kappa$. In all denoised images, the step size was 0.05 and the number of iterations 40. A higher diffusion parameter value leads to higher noise suppression and potentially edge blurring.

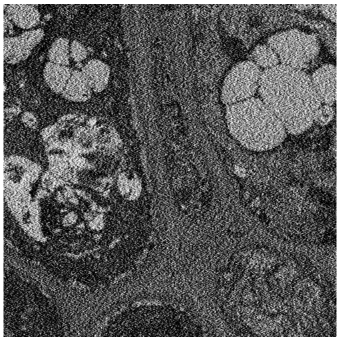

**(a)** Original noisy image

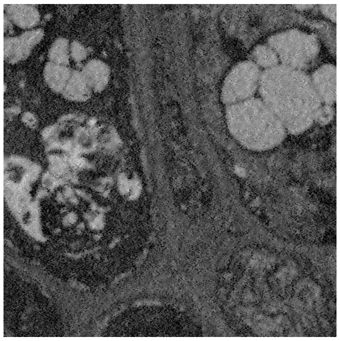

**(b)** $\sigma_{\text{int}} = 1$, $\sigma_{\text{sp}} = 3$

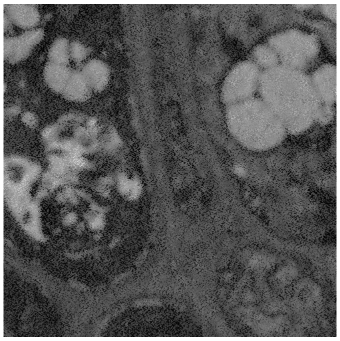

**(c)** $\sigma_{\text{int}} = 1$, $\sigma_{\text{sp}} = 5$

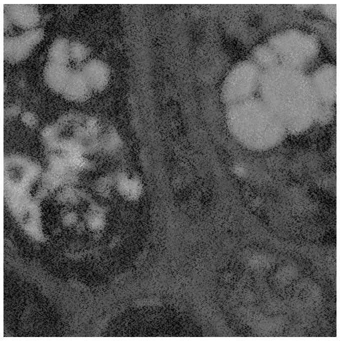

**(d)** $\sigma_{\text{int}} = 1$, $\sigma_{\text{sp}} = 7$

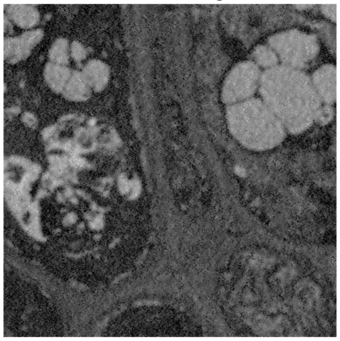

**(e)** $\sigma_{\text{int}} = 4$, $\sigma_{\text{sp}} = 3$

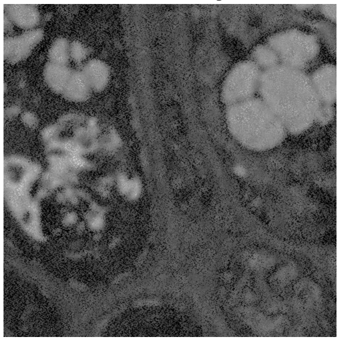

**(f)** $\sigma_{\text{int}} = 4$, $\sigma_{\text{sp}} = 5$

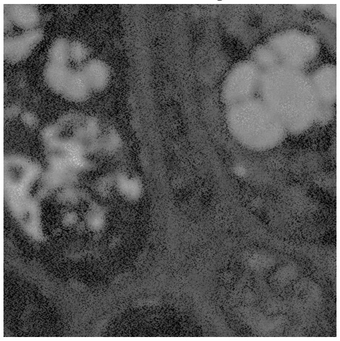

**(g)** $\sigma_{\text{int}} = 4$, $\sigma_{\text{sp}} = 7$

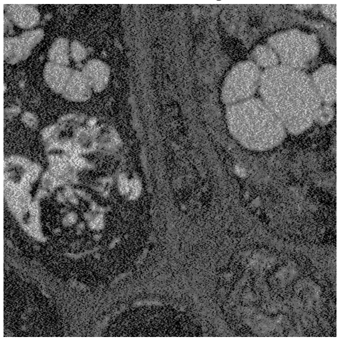

**(h)** $\sigma_{\text{int}} = 7$, $\sigma_{\text{sp}} = 3$

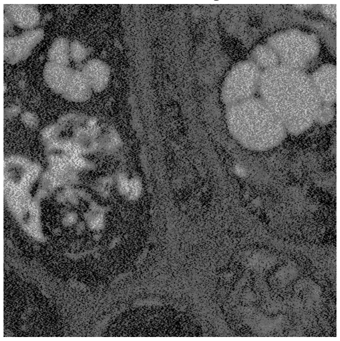

**(i)** $\sigma_{\text{int}} = 7$, $\sigma_{\text{sp}} = 5$

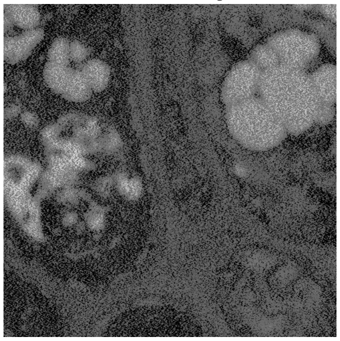

**(j)** $\sigma_{\text{int}} = 7$, $\sigma_{\text{sp}} = 7$

**Supplementary Figure 4.** Bilateral filtering. The top image is the original noisy image; the images underneath show the denoising behavior of the bilateral filter for 9 different combinations of spatial and intensity (range) damping factors: $\sigma_{\text{sp}} = 3, 5, 7$ and $\sigma_{\text{int}} = 1, 4, 7$. The parameter $\sigma_{\text{sp}}$ has the most significant influence on noise suppression, whereas $\sigma_{\text{int}}$ regularizes the intensity similarity between pixels that can be averaged.

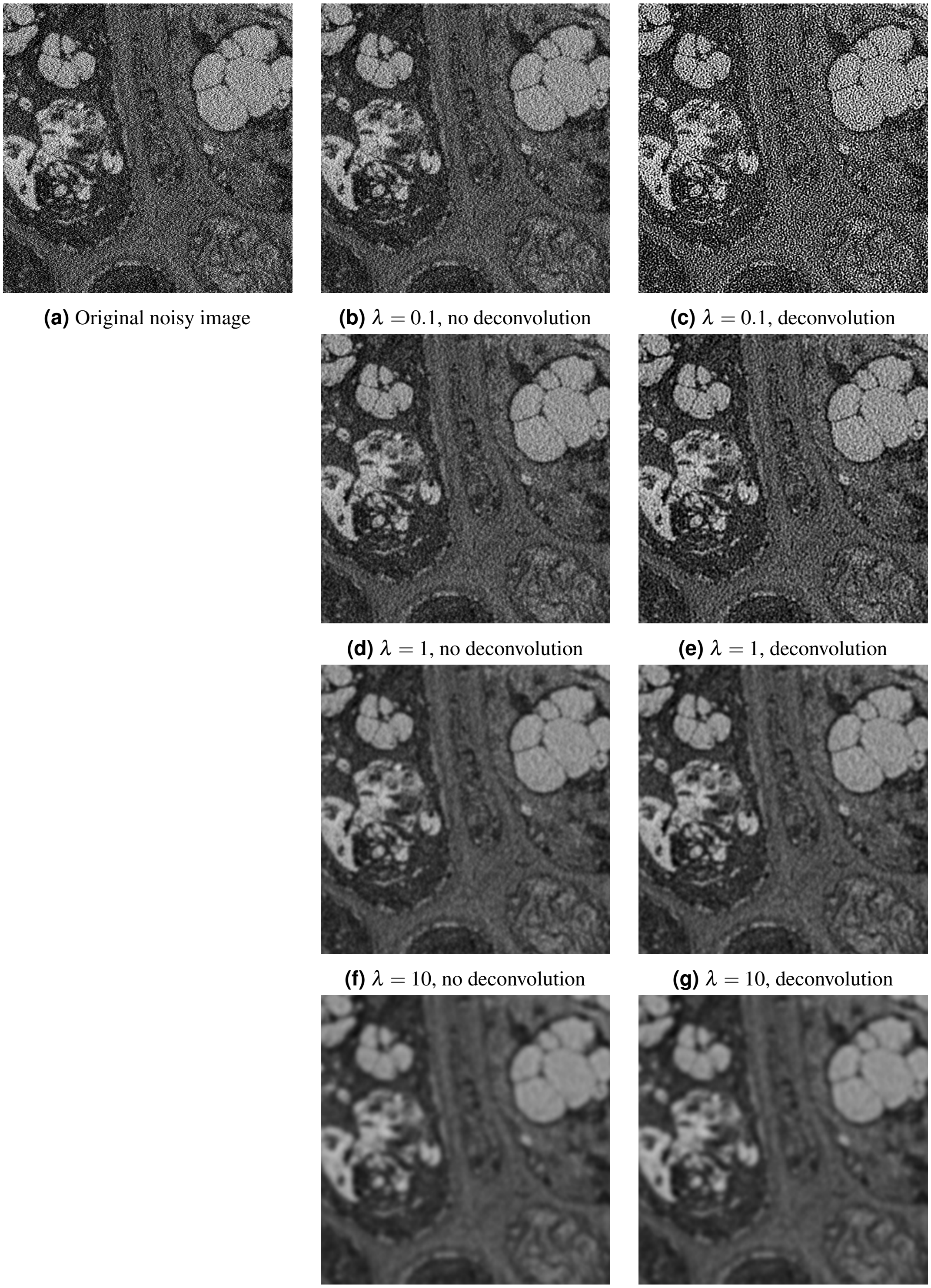

**(a)** Original noisy image **(b)** $\lambda = 0.1$, no deconvolution **(c)** $\lambda = 0.1$, deconvolution

**(d)** $\lambda = 1$, no deconvolution **(e)** $\lambda = 1$, deconvolution

**(f)** $\lambda = 10$, no deconvolution **(g)** $\lambda = 10$, deconvolution

**(h)** $\lambda = 100$, no deconvolution **(i)** $\lambda = 100$, deconvolution

**Supplementary Figure 5.** Tikhonov denoising, without deconvolution (middle column) and with deconvolution (right column). In all images the number of iterations was 100. For the images where deconvolution was applied, the blur kernel standard deviation had $\sigma = 1.5$. The $\lambda$ parameter offers control over the degree of noise suppression and deconvolution slightly sharpens the result.

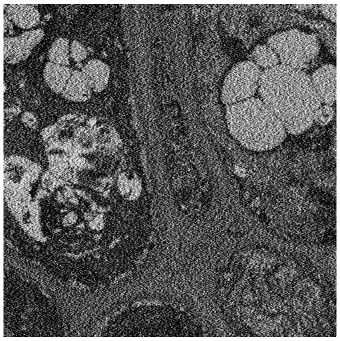

**(a)** Original noisy image

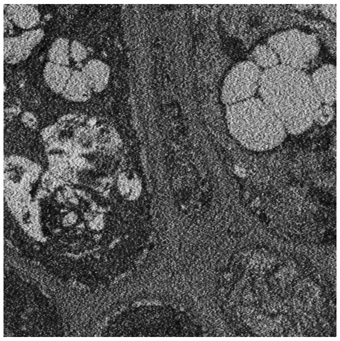

**(b)** $\lambda = 0.125$, 50 iterations

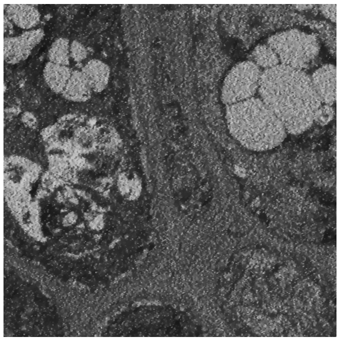

**(c)** $\lambda = 0.125$, 200 iterations

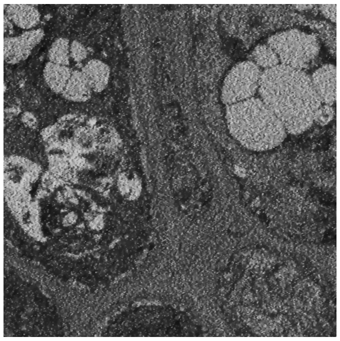

**(d)** $\lambda = 0.25$, 50 iterations

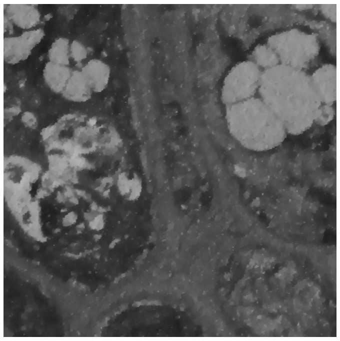

**(e)** $\lambda = 0.25$, 200 iterations

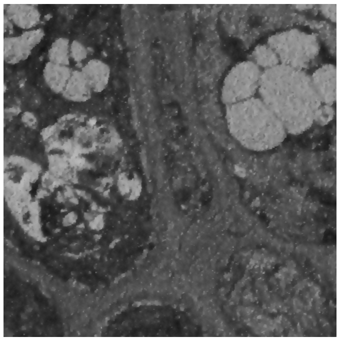

**(f)** $\lambda = 0.5$, 50 iterations

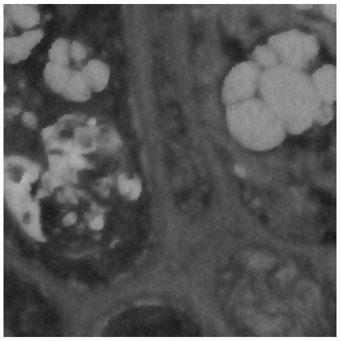

**(g)** $\lambda = 0.5$, 200 iterations

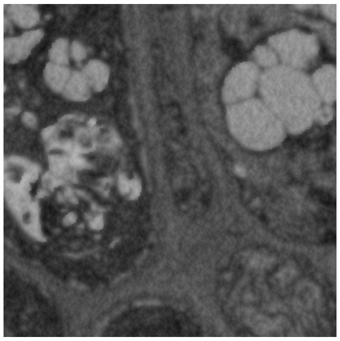

**(h)** $\lambda = 1$, 50 iterations

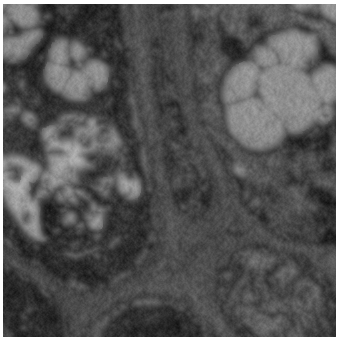

**(i)** $\lambda = 1$, 200 iterations

**Supplementary Figure 6.** Total Variation denoising. On the left is the original noisy image. From top to bottom are the denoised results for increasing values of the regularization parameter $\lambda$. The middle column used 50 iterations of the algorithm, the right column shows a more precise result using 200 iterations. The $\lambda$ parameter offers control over the degree of noise suppression and the amount of iterations is preferably as high as possible in order to guarantee convergence.

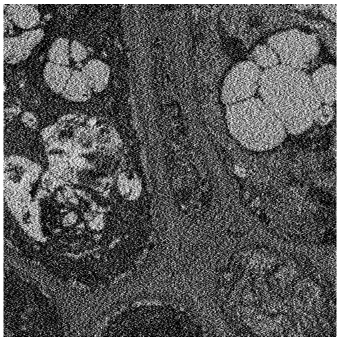

**(a)** Original noisy image

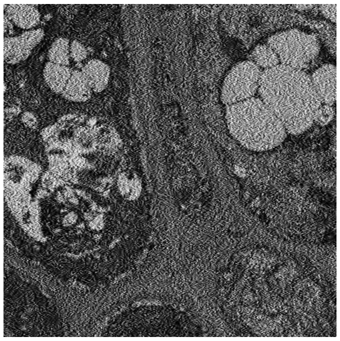

**(b)** $\sigma = 0.1, J = 3$

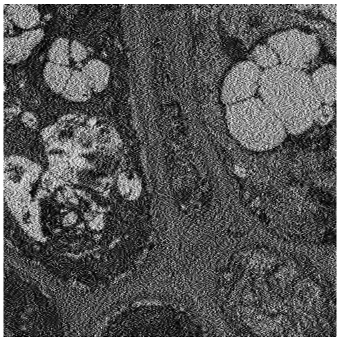

**(c)** $\sigma = 0.1, J = 4$

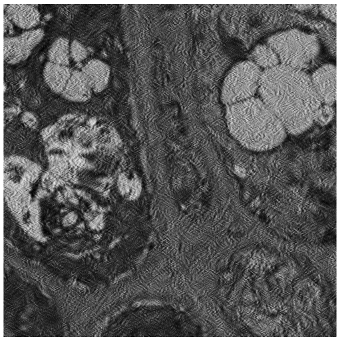

**(d)** $\sigma = 0.2, J = 3$

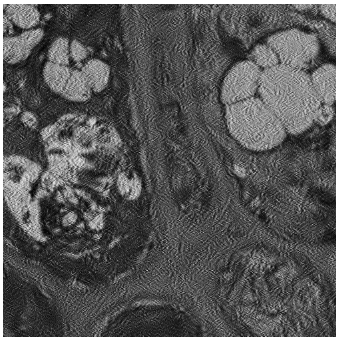

**(e)** $\sigma = 0.2, J = 4$

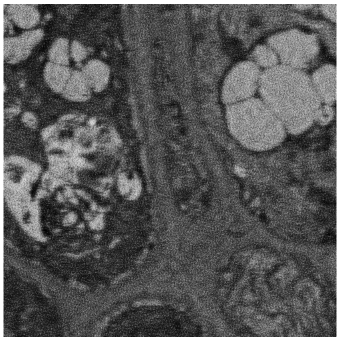

**(f)** $\sigma = 0.3, J = 3$

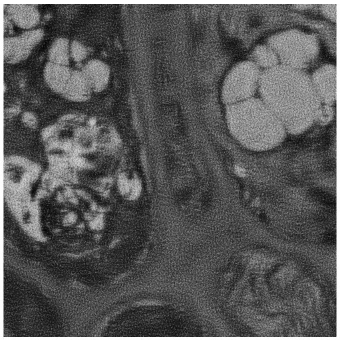

**(g)** $\sigma = 0.3, J = 4$

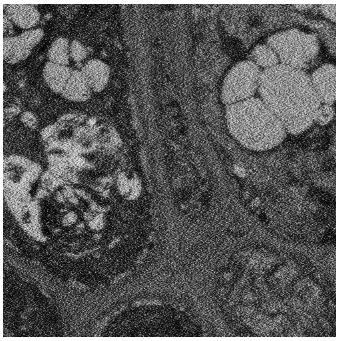

**(h)** $\sigma = 0.4, J = 3$

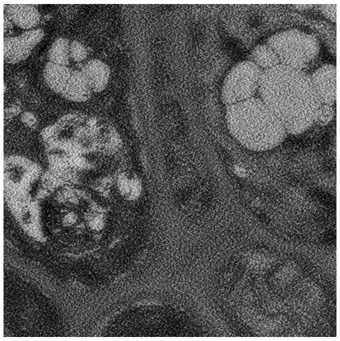

**(i)** $\sigma = 0.4, J = 4$

**Supplementary Figure 7.** BLS-GSM. The parameter $\sigma$ preferably matches the noise standard deviation, whereas an increasing number of scales typically leads to higher noise suppression.

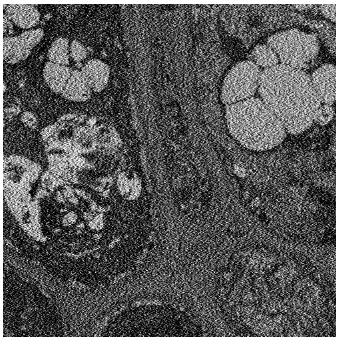

**(a)** Original noisy image

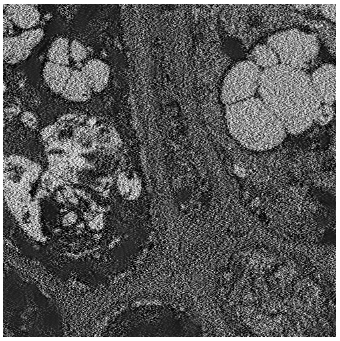

**(b)** $h = 0.35$, $W = 10$

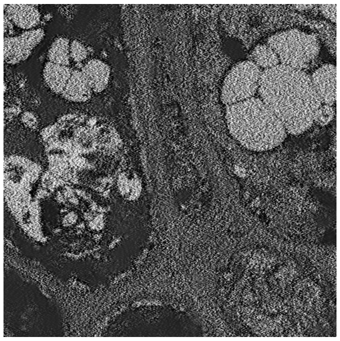

**(c)** $h = 0.35$, $W = 20$

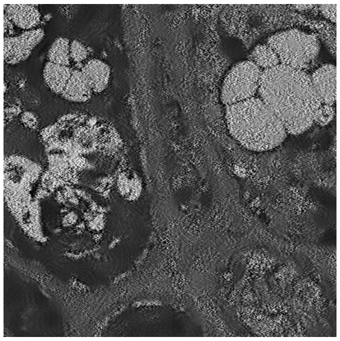

**(d)** $h = 0.4$, $W = 10$

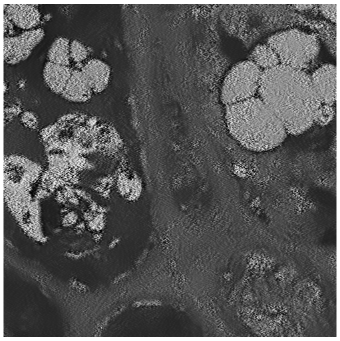

**(e)** $h = 0.4$, $W = 20$

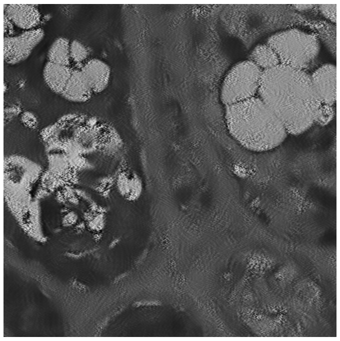

**(f)** $h = 0.45$, $W = 10$

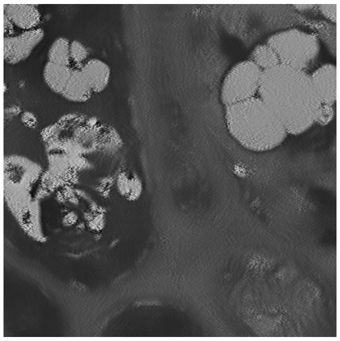

**(g)** $h = 0.45$, $W = 20$

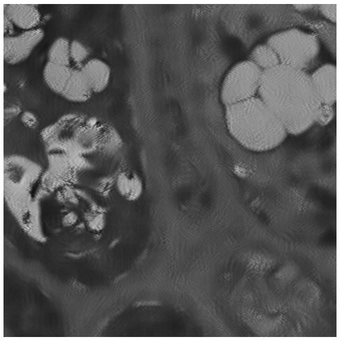

**(h)** $h = 0.5$, $W = 10$

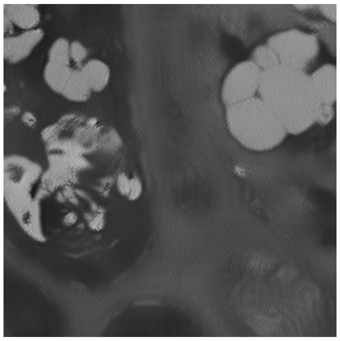

**(i)** $h = 0.5$, $W = 20$

**Supplementary Figure 8.** Non-local means denoising without deconvolution. The half block size used for averaging (similarity window) is $B = 7$ (*i.e.* a $15 \times 15$ window) in all images. Increasing the damping parameter $h$ leads to more noise suppression. Increasing the search window size leads to potentially better pixel candidates for averaging, but a higher computational cost.

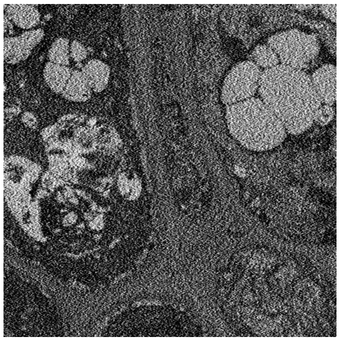

**(a)** Original noisy image

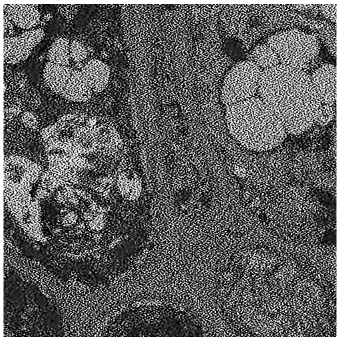

**(b)** $h = 0.35$, $\lambda = 0.5$

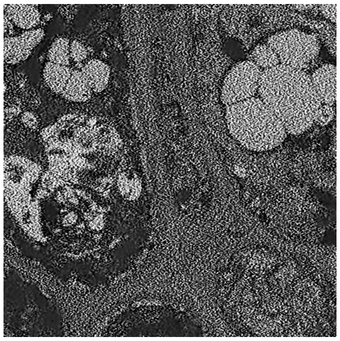

**(c)** $h = 0.35$, $\lambda = 2$

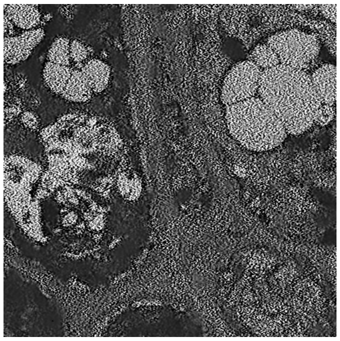

**(d)** $h = 0.4$, $\lambda = 0.5$

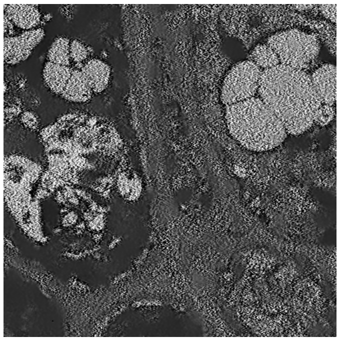

**(e)** $h = 0.4$, $\lambda = 2$

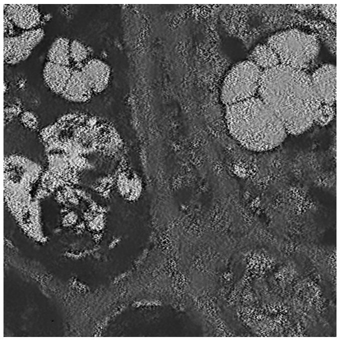

**(f)** $h = 0.45$, $\lambda = 0.5$

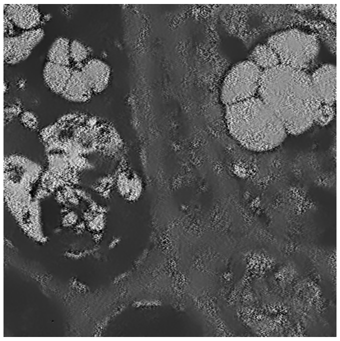

**(g)** $h = 0.45$, $\lambda = 2$

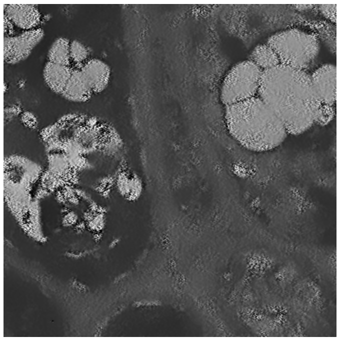

**(h)** $h = 0.5$, $\lambda = 0.5$

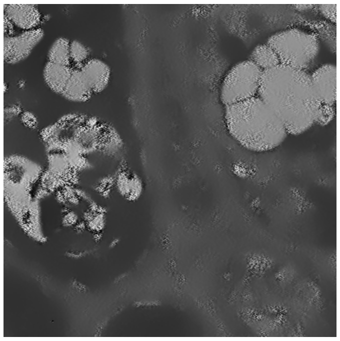

**(i)** $h = 0.5$, $\lambda = 2$

**Supplementary Figure 9.** Non-local means denoising with deconvolution. The half block size used for averaging (similarity window) is $B = 7$ (*i.e.* a $15 \times 15$ window) in all images. The half search window size is $W = 10$ (*i.e.* a $21 \times 21$ window) in all images. Lower values of the regularization parameter $\lambda$ emphasize deconvolution over denoising. Increasing the damping parameter $h$ leads to more noise suppression as opposed to sharpening.

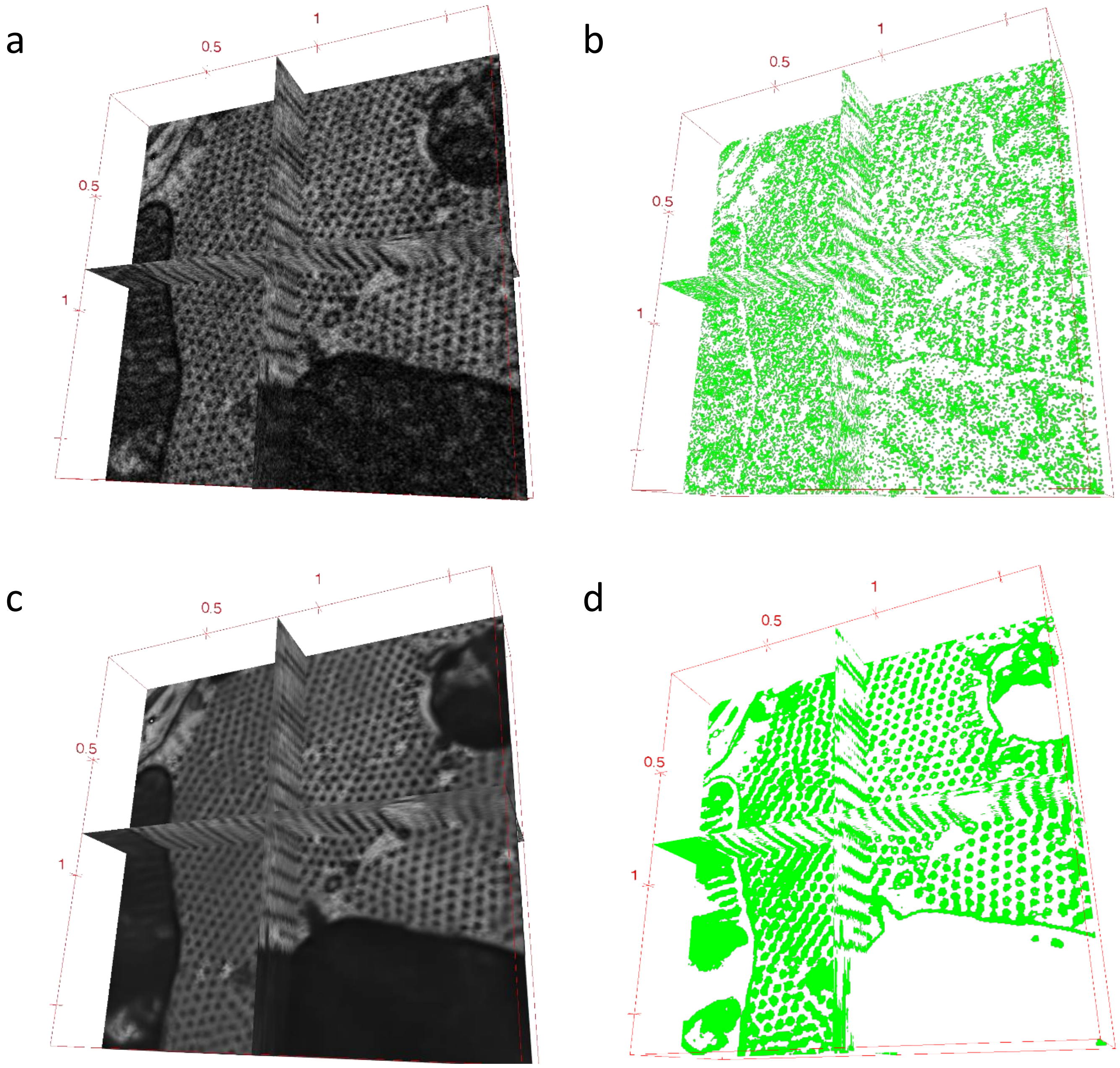


**Supplementary Figure 10.** (a) A 3D ROI from a FIB-SEM dataset was used for multi-orthogonal visualisation, using ImageJ 3D viewer. (b) A green mask created by applying intensity thresholding is shown in the same manner. (c) Similar visualisation was done on the same ROI, after denoising was applied. (d) Orthogonal views of the denoised data and mask after thresholding of the denoised data is shown.

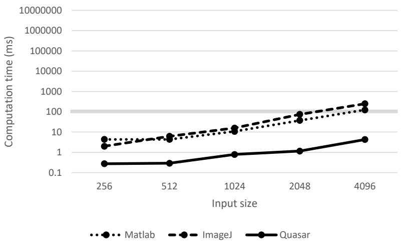


(a) Gaussian filter

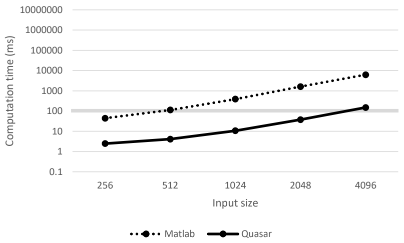


(b) Wavelet thresholding

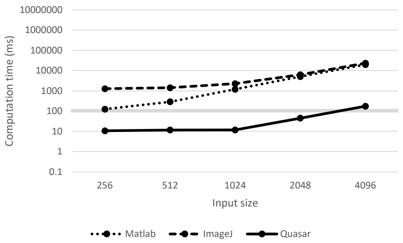


(c) Anisotropic diffusion

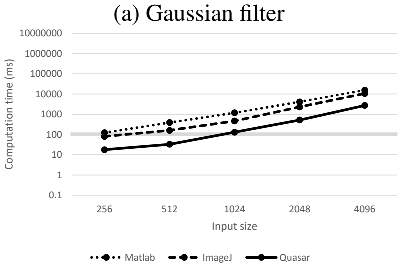


(d) Bilateral filter

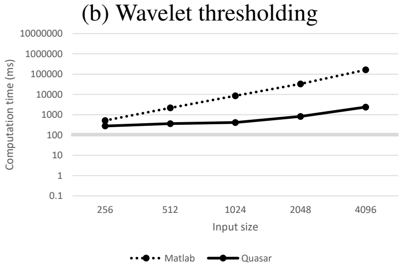


(e) BLS-GSM

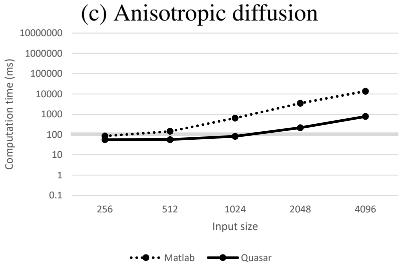


(f) Tikhonov denoising

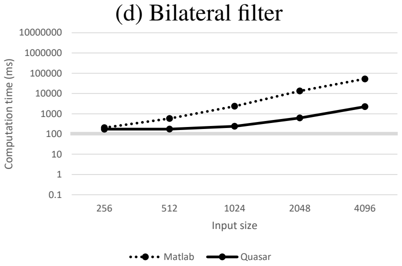


(g) Total variation denoising

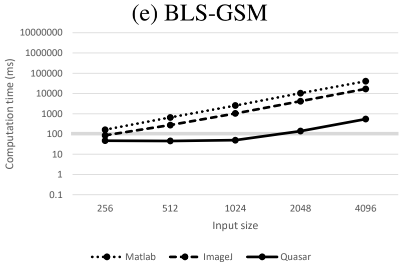


(h) Non-local means denoising

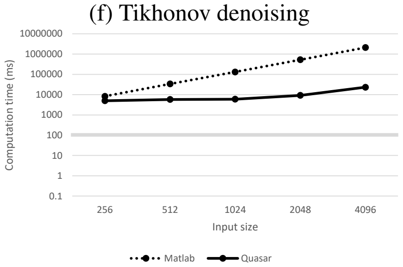


(i) Non-local means deconvolution

**Supplementary Figure 11.** Computational performance (in milliseconds) of denoising algorithms, as indicated, for different input sizes. A comparison is made between the proposed GPU-based Quasar framework and alternative CPU-based implementations in ImageJ[1–4] and MATLAB[5–13]. For each algorithm, we consider inputs of $256^2$, $512^2$, $1024^2$, $2048^2$ and $4096^2$ pixels. In general, the Quasar implementation performs one to two orders of magnitude faster compared to the existing software packages. 10 Hz is put as cut-off for real-time performance and indicated with a full line on the graph. Timings were measured using an Intel(R) Core(TM) i7-4930K CPU @ 3.40GHz and NVIDIA GTX 1070 GPU.
